# Molecular control of endurance training adaptation in mouse skeletal muscle

**DOI:** 10.1101/2023.02.18.529055

**Authors:** Regula Furrer, Barbara Heim, Svenia Schmid, Karl J.V. Nordström, Sedat Dilbaz, Danilo Ritz, Volkan Adak, Stefan A. Steurer, Jörn Walter, Christoph Handschin

## Abstract

Skeletal muscle has an enormous plastic potential to adapt to various external and internal perturbations. While morphological changes in endurance-trained muscles are well-described, the molecular underpinnings of training adaptation are poorly understood. We aimed at defining the molecular signature of a trained muscle and unraveling the training status-dependent responses to an acute bout of exercise. Our results reveal that even though at baseline, the transcriptomes of trained and untrained muscles are very similar, training status substantially affects the transcriptional response to an acute challenge, both quantitatively and qualitatively, in part mediated by epigenetic modifications. Second, proteomic changes were elicited by different transcriptional modalities. Finally, transiently activated factors such as the peroxisome proliferator-activated receptor γ coactivator 1α (PGC-1α) are indispensable for normal training adaptation. Together, these results provide a molecular framework of the temporal and training status-dependent exercise response that defines muscle plasticity in training.

**HIGHLIGHTS:**

1. Very few persistent transcriptional events define the trained muscle.
2. The training status determines the acute exercise response.
3. Epigenetic changes shape the transcriptional specification of trained muscle.
4. Absence of the key regulator PGC-1α causes suboptimal training adaptations.

## INTRODUCTION

Skeletal muscle exerts pleiotropic functions, from thermoregulation through shivering, to endocrine signaling by myokines and myometabolites, and detoxification of endogenous compounds, e.g. kynurenines or aberrantly high levels of ketone bodies^1–5^. However, the main task of skeletal muscle is the generation of force for different types of contractile activities, including strength, endurance, fine motor control, posture or breathing. Skeletal muscle thus not only exhibits broad morphological and functional specification, but also a remarkable adaptive plasticity to react to internal and external perturbations^4^. Remodeling of skeletal muscle requires interventions that disrupt homeostasis, to which muscle will progressively adapt only if repeated over time^3,4^. Morphologically, such adaptations are well described, e.g. fiber hypertrophy in resistance, or mitochondrial expansion, vascularization and energy substrate storage in endurance training^4^. In light of the powerful health benefits conferred by either modality of exercise^6,7^, it however is surprising that the molecular underpinnings of muscle plasticity in exercise are still only rudimentarily understood^4^. In particular, the mechanistic framework that links the perturbations evoked by individual, acute exercise bouts to long-term training adaptations are largely unknown^3,4^. Additionally, it is unclear how the training status affects the molecular response to an acute bout of exercise, and how changes in gene expression are ultimately linked to persistent modulation of protein levels, organelle function and tissue plasticity. Functionally, a repeated bout effect has been postulated based on the observation of reduced muscle damage and soreness in trained compared to naïve muscle^8,9^. Accordingly, a diminished amplitude in the expression of a number of genes in repeated exercise bouts has been reported, at least with constant training load^10,11^, nevertheless resulting in steady accumulation of transcripts, proteins and performance over time^12–17^. Training habituation and attenuation of the respective responses could thus define this process, with increasingly higher intrinsic resilience and lower perturbations of muscle cells. Such an encompassing model of transcriptional attenuation in training adaptation however is contradicted by different observations. For example, a broad-ranging qualitative and quantitative specification is implied by the vastly different epigenetic modifications in acute and chronic exercise settings^17–19^. Accordingly, the expression of many genes is not following an attenuating pattern, instead showing an exacerbated response in trained muscle, as described for the peroxisome proliferator-activated receptor γ coactivator 1α (PGC-1α)^20^. Of note, PGC-1α and most other postulated key regulatory factors in exercise adaptation in skeletal muscle, including the AMP-dependent protein kinase (AMPK), the mammalian target of rapamycin (mTOR), or the calcium-dependent proteins calcineurin A (CnA) and calcium/calmodulin-dependent kinase (CaMK) are only very transiently modulated^4^. Therefore, little knowledge about the chronic, persistent mechanistic network in training adaptation exists. To understand these fundamental aspects of muscle biology and plasticity, we therefore studied the acute endurance exercise and chronic training response of mouse muscle in a systematic and comprehensive manner. Based on the interrogation of the molecular underpinnings of epigenetic, transcriptional and proteomic changes, we provide a novel mechanistic framework of endurance training adaptations. The transcriptomic data of acute exercise and chronic training are provided in the myo-transcriptome of exercise database (myoTrEx, *LINK*).

## RESULTS

### The transcriptome of a trained is surprisingly similar to that of an untrained muscle

To study differences between naïve and endurance-trained muscles, mice were exercised by treadmill running on 5 days per week for 1 hour (Figure S1A). After 4 weeks, a significant improvement in running performance was observed (Figure S1B). A proteomic analysis also indicated a substantial remodeling of skeletal muscle (Figure 1A; Table S1). For example, proteins involved in mitochondrial respiration, lipid metabolism, oxygen transport, or stress resilience are more abundant in trained than untrained muscle (Figures 1B, 1C, S1C, S1D; Table S2). In contrast, the levels of proteins linked to catabolic processes related to proteasomal degradation are mitigated by endurance training (Figures 1B & S1E), which, together with the induced molecular chaperones, alludes to altered proteostasis. According to prevailing models, these proteomic changes are brought about by a persistent modulation of gene expression with repeated exercise bouts^13^. We therefore assessed the transcriptomic landscape of the trained muscle. Interestingly, only a small number of genes (<2% of the detected genes) was significantly altered (Figure 1D). These few genes define long-term cellular changes, e.g. related to fiber-type switch, metabolic remodeling or decreased inflammation (Figure 1E; Table S3). In line with these observations, Integrated System for Motif Activity Response Analysis (ISMARA)^21^ revealed higher activity of the Esrrb_Esrra and lower activity of Rela_Rel_Nfkb1 motifs (Figure 1F; Table S4). Surprisingly, the altered proteome of a trained muscle is only to a small extent maintained transcriptionally. These proteins are predominantly involved in lipid metabolic process (Figure S1F; Table S5). Thus, the majority of the proteins that define the long-term plasticity of a trained muscle is not directly linked to a corresponding transcriptional response.

**Figure 1.**
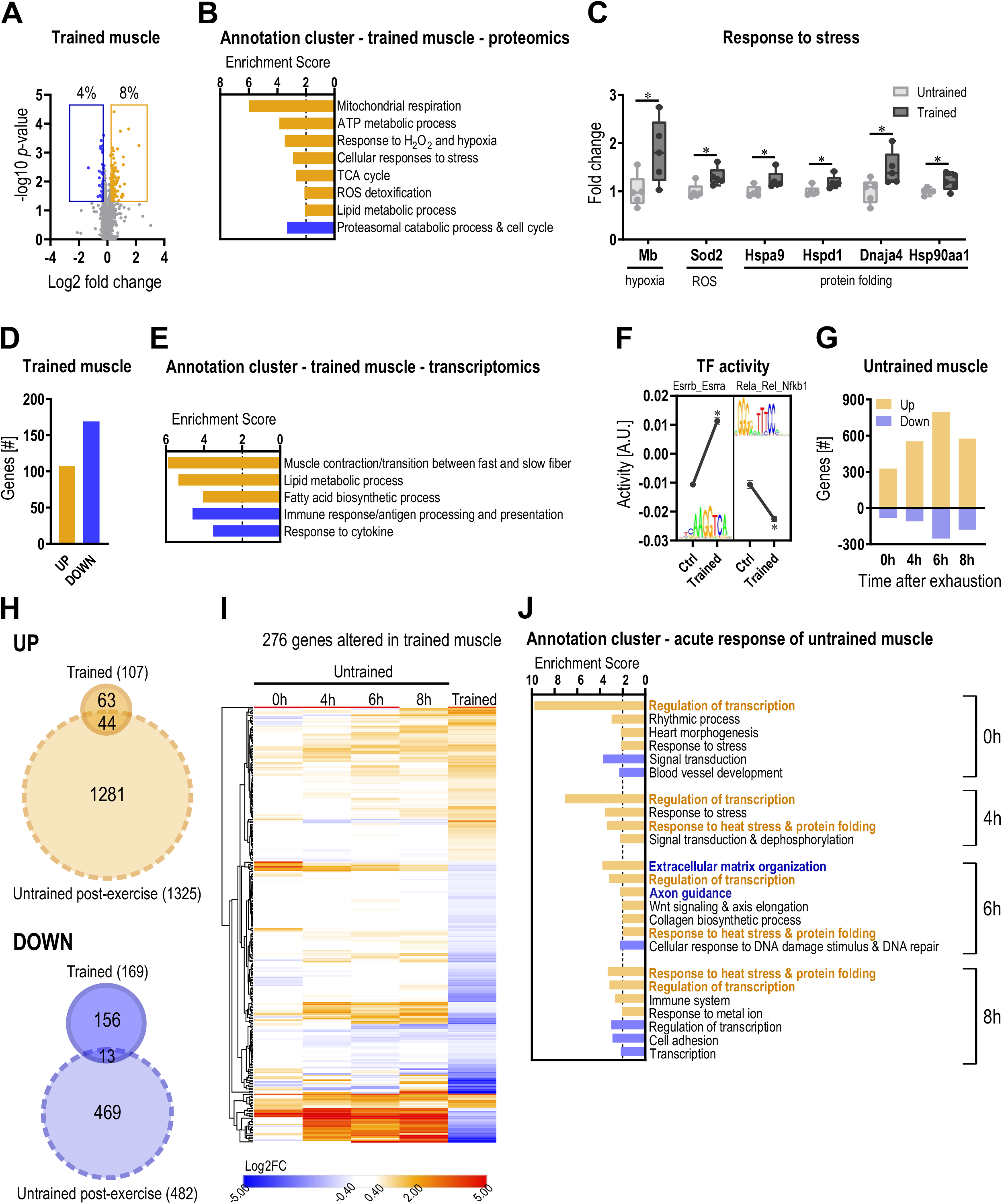
The transcriptome of a trained is similar to that of an untrained muscle. (A) Volcano plot of all detected proteins in trained muscle using mass spectrometry-based proteomics (orange = higher abundant; blue = lower abundant; cutoff: *p* <0.05; Log2FC ±0.2). (B) All functional annotation clusters of up- (orange) and downregulated (blue) proteins in trained muscle with an enrichment score >2. (C) Examples of proteins involved in the response to stress in sedentary untrained (light gray) and unperturbed training (dark gray) muscle. (D) Number of genes differentially expressed in unperturbed trained muscle (cutoff: FDR <0.05; Log2FC ±0.6). (E) All functional annotation clusters of up- (orange) and downregulated (blue) genes in trained muscle with an enrichment score >2. (F) Motifs of transcription factors from ISMARA that are among those with the highest and lowest activity. (G) Number of genes after an acute bout of exhaustion exercise that are up- (orange) and downregulated (blue). (H) Venn diagram of all genes that are regulated after an acute bout of exercise (light color, dashed line) and those that are changed in unperturbed trained muscle (orange = upregulated; blue = downregulated). (I) Heatmap of all genes differentially expressed in unperturbed trained muscle to visualize the overlap with acutely regulated genes using euclidean distance hierarchical clustering for rows. (J) All functional annotation clusters of up- (orange) and downregulated (blue) genes after an acute exercise bout in untrained muscle with an enrichment score >2. Data from 5 biological replicates. Data represent means ± SEM. Statistics were performed using empirical Bayes moderated t-statistics for proteomics and within the CLC genomics workbench software for RNAseq data. * indicates difference to Ctrl (pre-exercise condition) if not otherwise indicated; in (C) * *p*-value <0.05; in (F) * z-score >1.96. See also Figure S1; Tables S1-S4.

These unexpected results, defining the endurance-trained muscle as a largely non-transcriptional event, raised the question whether perturbations evoked by acute events activate transcriptional networks that encode the biological programs observed in trained muscle. To test this hypothesis, training-naïve mice were exercised to exhaustion by treadmill running, and the muscle transcriptome was assessed 0h, 4h, 6h and 8h post-exhaustion (Figure S1A). Similar to other studies, we found a large number of gene regulatory events in this context, peaking 6h post-exhaustion (Figure 1G). A subset of these acute changes correlated with the accumulation of proteins in a trained muscle. These proteins were mostly upregulated, and predominantly involved in aerobic respiration (Figure S1G; Table S5). Intriguingly, the acutely regulated genes only poorly overlapped with persistent transcriptomic changes in trained muscle, as only 21% (57 of the 276) of the genes modulated in an unperturbed trained muscle are also regulated acutely in naïve muscle (Figures 1H, 1I, S1H-L). In fact, some of the genes exhibited opposite regulation (Figures 1I, S1M), e.g. reflected in transcripts related to inflammation (up acutely post-exercise, down in trained muscle). Functionally, many of the acutely regulated genes were related to a strong transcriptional response, and to various aspects of stress response, damage, axon guidance, and extracellular matrix (ECM) organization (Figure 1J; Table S3).

### Qualitative and quantitative differences in the transcriptional response to an acute bout of exercise of a trained and untrained muscle

Since the acute response in naïve muscle was not predictive of training changes, we next investigated the response of a trained muscle to an acute bout of endurance exercise at the same four time points. Accordingly, mice that were trained for 4 weeks performed an exhaustive bout of treadmill running (Figure S1A). Most strikingly, the transcriptomic responses of naïve and trained muscle to an acute endurance exercise bout were decisively different, qualitatively and quantitatively, the latter both in terms of amplitude (extent of change) and phase (temporal regulation) (Figures 2A-C). First, less than half of the upregulated genes overlapped between these two conditions, and even a smaller proportion of the downregulated transcripts, of which a greater number was altered in the trained condition (Figure 2B). Strikingly, the functional prediction of the acute response of trained muscle was diametrically opposite to that of the untrained in regard to ECM remodeling and axon guidance (Figures 1J, 2C, S2A-D; Table S3). Many of these predicted functions, including a modulation of inflammation, could originate from non-myocytes in muscle tissue. Therefore, we performed cellular deconvolution of the bulk with published single cell and single nucleus RNAseq data of untrained muscle (Figure 2D)^22,23^. These analyses predict a surprisingly detailed specification of gene expression between different cell types (Figures 2E-G, S2E, S2F). For example, while ECM remodeling is mainly driven by fibro-adipogenic progenitors (FAPs) in untrained muscle, tenocytes are more involved in the adaptive processes of ECM in trained muscle. These cell type-specific responses presumably result in the corresponding distinct outcomes for ECM remodeling, axon guidance, and potentially other functions after an acute endurance exercise bout in naïve compared to trained muscle.

**Figure 2.**
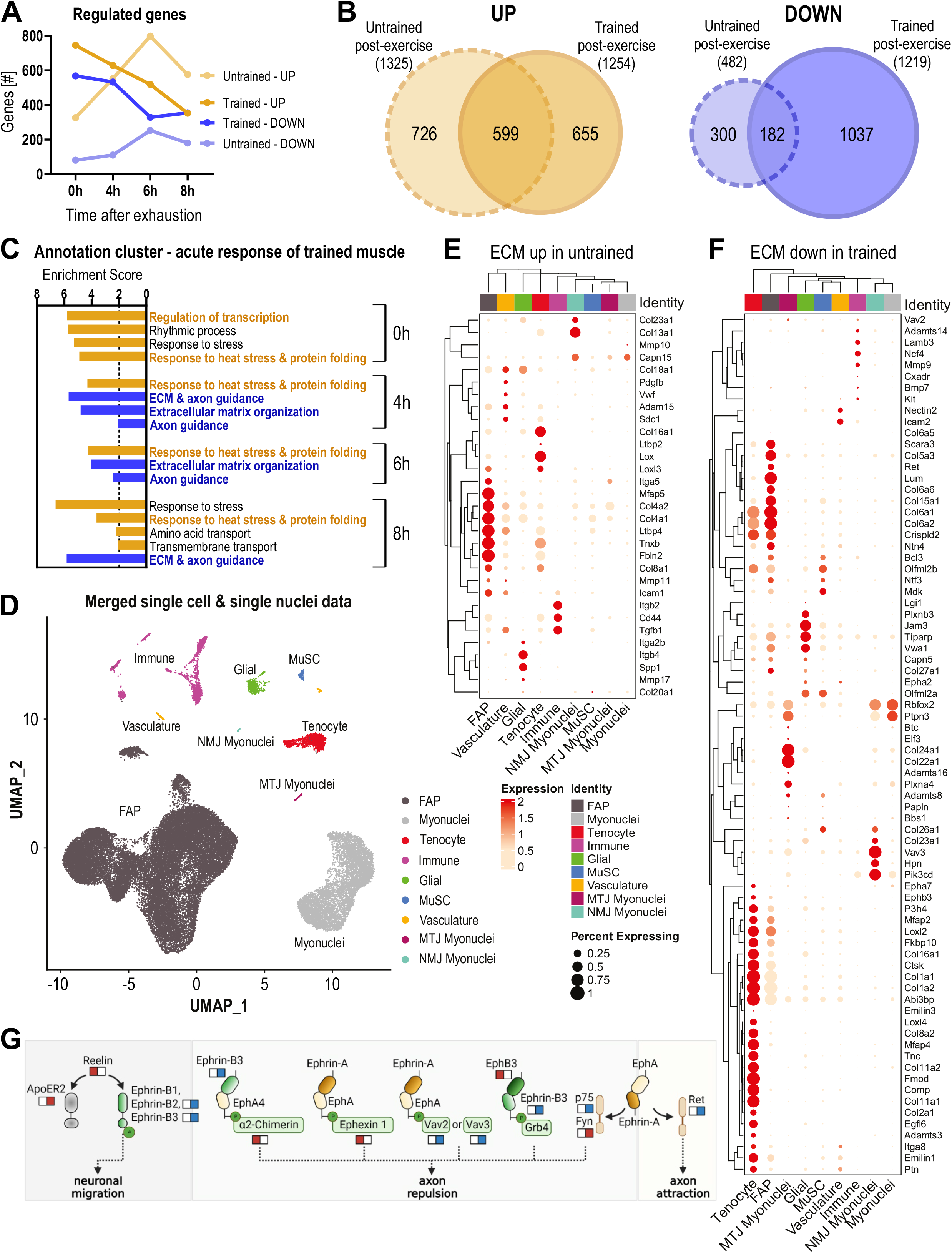
Qualitative transcriptional response to exercise depends on training status. (A) Number of genes differentially expressed immediately (0h), 4h, 6h and 8h after an acute bout of exhaustion exercise (cutoff: FDR <0.05; Log2FC ±0.6) in untrained and trained muscle. (B) Venn diagram of all significantly up- (orange) and downregulated (blue) genes (all time points merged) in untrained (light color, dashed line) and trained (dark color, solid line) muscle. (C) All functional annotation clusters of up- (orange) and downregulated (blue) genes after an acute exercise bout in trained muscle with an enrichment score >2. (D) UMAP plot of public available single cell and single nucleus RNAseq datasets^22^’^23^ to demonstrate the cellular specification of the exercise response in muscle (FAP = fibro-adipogenic progenitors; MuSC = muscle stem cells; MTJ = myotendinous junction; NMJ = neuromuscular junction). (E-F) Deconvolution of genes involved in ECM remodeling that are upregulated in untrained (E) and downregulated in trained (F) muscle. (G) Schematic representation of genes involved in axon guidance and the possible functional consequences^50^. The left square below each gene name represents the untrained response and the right square the trained response. Red = upregulated; blue = downregulated (illustration was created with BioRender.com). Data from 5 biological replicates. Data represent means ± SEM. Statistics of RNAseq data were performed within the CLC genomics workbench software. * indicates difference to Ctrl (pre-exercise condition); *<0.05; **<0.01; ***<0.001. See also Figure S2; Table S3.

In contrast to the qualitative differences in functional gene clusters between untrained and trained muscle, common processes induced upon an acute perturbation such as the regulation of transcription or response to heat stress also exhibit a strong training status-dependent specificity (Figures 1J, 2C). For example, qualitative differences in the regulation of transcription factors such as early growth response 3 (Egr3), or quantitative modulation of transcriptional regulators such as PGC-1α were observed (Figure 3A). ISMARA confirmed the substantial regulatory diversification (Figure S3A-F; Table S4). While approximately 35-43% of the motifs are specific to the training status (Figure S3B), many of the common motifs (n=77) show altered trajectories and/or amplitudes (Figures S3C-F). Of note, despite the >50% overlap between motif activities in untrained and trained muscle, in 21 of the 77 motifs, the activity profiles point in the opposite direction. For example, the Wrnip1_Mta3_Rcor1 motif activity is higher in untrained and lower in trained muscle and, based on the association with collagen formation, could contribute to the distinct patterns of ECM remodeling (Figure 3B).

**Figure 3.**
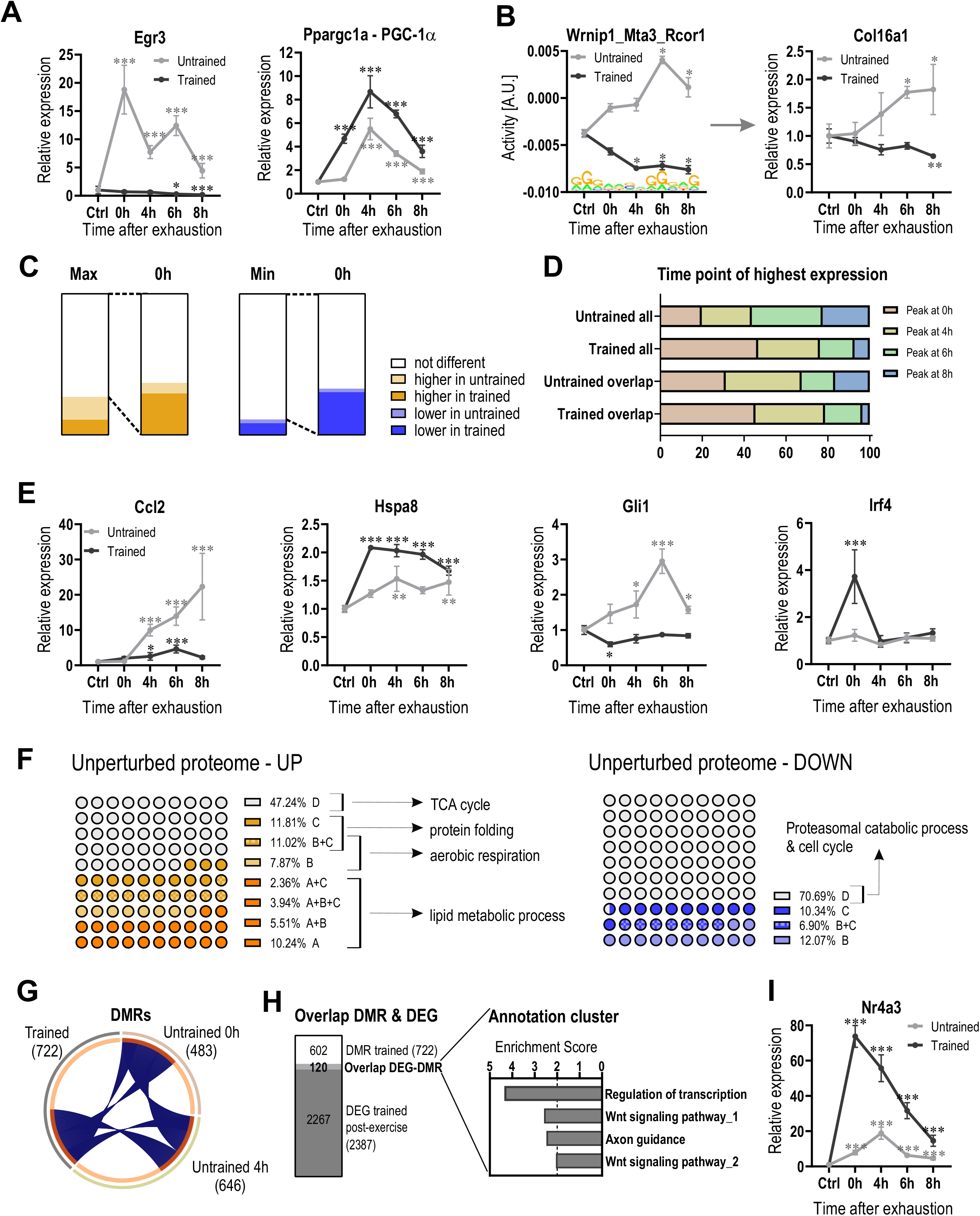
Faster transcriptional response in trained muscle after one bout of exhaustion exercise. (A) Examples of distinct gene signatures represented in the annotation cluster “regulation of transcription” in untrained (light gray) and trained (dark gray) muscle. (B) Motif activities from ISMARA and expression changes of a predicted target gene that show an opposite regulation in untrained and trained muscle. (C) Proportion of overlapping genes with either the same maximal/minimal fold change (white), higher amplitude (orange = upregulated; blue = downregulated) in untrained muscle (light color) or higher amplitude in trained muscle (dark color). The second bar graph (named 0h) represents the proportion of genes with the same maximal value and depicts the proportion of gene that have the same fold change at the 0h time point (white), higher amplitude in untrained muscle (light color) or higher amplitude in trained muscle (dark color). (D) Representation of the time points when peak expression is reach in either all upregulated genes (upper 2 bars) or only the 599 commonly upregulated genes (lower 2 bars). (E) Examples of different gene trajectories in untrained and trained muscle after an acute exercise bout representing the different training status-specific transcriptional scenarios. (F) Relative transcriptional contribution to the proteome of an unperturbed trained muscle: A/dark orange = mRNA unperturbed trained muscle, B/light orange/light blue = mRNA acute response untrained muscle, C/orange/blue = mRNA acute response trained muscle, D/white = protein only (unperturbed trained muscle). (G) Circos plot of differentially methylated regions (DMRs) 0h, 4h post-exercise and in unperturbed trained muscle. (H) Bar Venn diagram of DMRs of an unperturbed trained muscle (white) and differentially expressed genes (DEG) after acute exercise in trained muscle (dark gray) and the functional annotation clusters of the overlap (light gray, n = 120) with an enrichment score >2. (I) Example of a transcription factor that is differentially methylated in trained muscle and higher expressed after exercise in trained (dark gray) compared to untrained muscle (light gray). Data from 5 biological replicates. Data represent means ± SEM. Statistics were performed using empirical Bayes moderated t-statistics for proteomics and within the CLC genomics workbench software for RNAseq data. Difference in relative expression changes presented in (D) were calculated using a two-tailed Student’s t-test. * indicates difference to Ctrl (preexercise condition); *<0.05 (for motif activity: * z-score >1.96); **<0.01; ***<0.001. See also Figures S3, S4; Tables S4-S6.

Intriguingly, even the genes with converging expression patterns in untrained and trained muscle often exhibit quantitative differences. While the maximal and minimal amplitude of most commonly regulated genes is very similar in untrained and trained muscle, a substantially higher proportion of these genes display a higher amplitude in trained muscle 0h post-exercise (Figures 3C, S4A). In fact, the peak expression of many of the overlapping genes shifted towards earlier time points in the trained compared to the untrained muscle (Figures 3D, S4B, S4C). Accordingly, almost half of all upregulated genes in trained muscle peak at 0h while this only applies to ~20% of the upregulated genes in untrained muscle (majority peaks after 6h) (Figure 3D). Overall, opposed to the model of general attenuation of gene expression with training habituation^3,10,11,13–15^, our results suggest a much more complex picture, with significant occurrence of all scenarios: attenuation, exacerbation, and selective expression changes in the naïve or trained muscle after an acute exercise bout (Figure 3E). Moreover, most of the acute transcriptional changes are not retained in an unperturbed trained muscle (Figure S4D). However, collectively, these gene regulatory events correlate with the trained proteome and explain up to 43% for upregulated, and 30% for downregulated proteins for which transcript data are available (Figure 3F), highlighting the importance of broad comparisons between transcriptomes and proteomes^24^. Interestingly, proteins that are transcriptionally sustained in a trained muscle (22% of the proteins) are involved in lipid metabolic process (Figure 3F; Table S5). In contrast, genes that are transiently induced upon acute exercise in untrained or trained muscle contribute to the elevation of proteins involved in aerobic respiration and protein folding, respectively. The 47% of the proteome for which no transcriptional regulation was seen could either be underlying transcriptional events at different time points than analyzed in this study, or primarily be affected by post-transcriptional mechanisms. Interestingly, many of these proteins cluster in the tricarboxylic acid (TCA) cycle. Overall, these findings allude to a complex regulatory network by which long-term adaptations are brought about.

### Priming of regulatory genes in trained muscle by DNA methylation changes

Next, we studied mechanistic processes that contribute to the divergent specification of exercise-induced gene expression in naïve and trained muscle. Epigenetic changes have previously been reported in training adaptation^17,19^. We therefore performed reduced representation bisulfite sequencing (RRBS) to catalogue DNA methylation events in the trained muscle and those elicited by acute exercise. An acute endurance exercise bout resulted in a number of DNA methylation changes at 0h and 4h, some of which could be associated with gene expression changes (Figures S4E, S4F). Such transient epigenetic regulation of transcription has been described for PGC-1α and other exercise-responsive genes^18^. Intriguingly, the chronically retained DNA methylation events in trained muscle differed substantially from these acute changes (Figure 3G, S4G). Of note, almost none of these differentially methylated regions (DMRs) could be associated with the gene expression changes in trained muscle (Figure S4H). In contrast, a subset of these DMRs are in the immediate genomic vicinity of a small subset of genes (120 out of 2387) that are regulated after an acute exercise bout in the trained muscle (Figure 3H), indicating that these epigenetic modulations could contribute to a priming of transcriptional regulation in this context. Intriguingly, these genes enrich in functions related to the regulation of transcription, Wnt signaling and axon guidance signaling effectors (Figures 3H, S4I-K; Table S6). For example, the exercise induction of nuclear receptor 4A3 (Nr4a3), which is associated with DMRs in the trained muscle, is not only greatly exacerbated in the trained compared to the untrained muscle, but also displaying a phase-shift with a peak immediately post-exercise (Figure 3I). Thus, epigenetic modifications could contribute to the different gene expression of a trained muscle upon an acute perturbation, primarily affecting regulatory genes, with subsequent downstream consequences independent of DNA methylation changes.

### PGC-1α is indispensable for a normal transcriptional response to acute exercise and long-term training

Notably, many transcriptional regulators that are engaged strongly and early after acute exercise exhibit a diversification between the first and, in our setting, 22^nd^ bout, thus naïve and trained muscle, including PGC-1α (Figure 3A). This coregulatory protein has been strongly implicated in the acute response by integrating various signaling pathways, and subsequently affecting the activity of numerous transcription factors, thereby coordinating a complex transcriptional network^25^. Our observation, recapitulating prior results in human muscle^20^, of a quantitative difference of PGC-1α in trained compared to untrained muscle would indicate that PGC-1α not only controls an acute stress response, but might also affect the transcriptome of exercised muscle in the trained state. However, the relevance of adequate regulation and function of PGC-1α in long-term training adaptations has not been established, and at least in part conflicting findings have been reported^26–29^. To obtain comprehensive information on muscle PGC-1α in training, we therefore repeated the exercise study with muscle-specific PGC-1α knockout (mKO) mice. In agreement with previous work^30^, mKO mice exhibit a reduced endurance capacity, running approximately 40% less than WT controls (Figure 4A). Nevertheless, the PGC-1α loss-of-function animals substantially improved maximal performance after 4 weeks of training, in relative and absolute terms, however still significantly less than the WT counterparts (Figures 4A, 4B). Moreover, maximal oxygen consumption (VO_2max_) failed to improve in mKOs (Figure 4C), alluding to an alternative adaptation of endurance capacity in these mice. In the acutely perturbed, training-naïve muscle, a massive blunting of transcriptional induction at the later time points (4-8h post-exercise) was found, thus subsequent to the physiological PGC-1α elevation in WT muscle (Figures 4D-J). Overall, absence of muscle PGC-1α affected 56% of all up-, and 65% of all downregulated exercise-responsive genes (Figure 4K). Some of these genes are functionally related to ECM remodeling, Wnt signaling and microglial cell proliferation (Figures 4L-N, S5A; Table S7), linked to a corresponding mitigation of Wrnip1_Mta3_Rcor1 and other transcription factor motifs related to the regulation of these processes (Figure S5B; Table S4). Genes which were not qualitatively affected by the absence of muscle PGC-1α contribute to transcriptional regulation and response to heat stress and protein folding (Figures 4G, 4J, S5C; Table S7). Intriguingly, the expression of a notable number of genes is only modulated in mKO muscles (Figure 5SC), for example a number of transcripts encoding proteins involved in inflammation, confirming prior descriptions of exacerbated activity-induced muscle damage and inflammation in mKO animals^30^.

**Figure 4.**
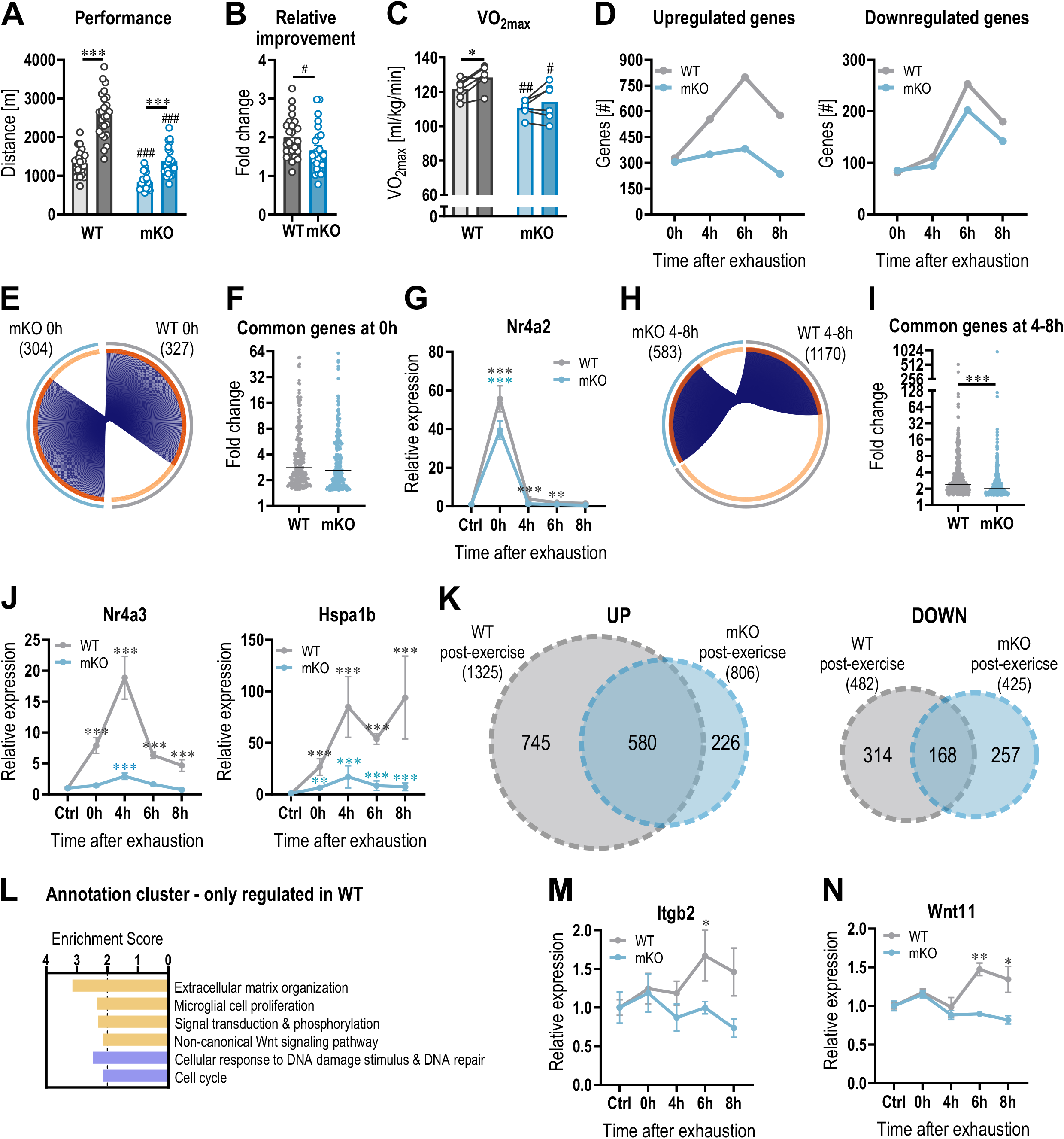
PGC-1α is indispensable for a normal transcriptional response to acute exercise and long-term training. (A) Performance of untrained (light color) and trained (dark color) WT (gray) and mKO (blue) animals. (B) Relative improvement of WT and mKO animals after 4 weeks of progressive treadmill training. (C) Changes in VO_2max_ before (light color) and after (dark color) training. (D) Number of genes that are up- and downregulated 0h, 4h, 6h and 8h after an acute exercise bout in untrained WT and mKO animals. (E) Circos plot of all upregulated genes immediately post-exercise (0h) in untrained WT and mKO mice. (F) Relative fold change (and median) of all commonly regulated genes at the 0h time point in untrained muscle. (G) Example of gene trajectories with the peak expression immediately post-exercise in untrained WT and mKO animals. (H) Circos plot of all upregulated genes after 4-8h (merged together) in untrained WT and mKO mice. (I) Relative fold change (and median) of all commonly regulated genes 4-8h post-exercise in untrained muscle. (J) Examples of gene trajectories with a peak at a later time point in untrained WT and mKO animals. (K) Venn diagrams of all up- and downregulated genes after an acute bout of exercise in untrained WT (light gray) and mKO (light blue) mice. (L) All functional annotation clusters of up- (orange) and downregulated (blue) genes that are only regulated in untrained WT mice (745 genes up- and 314 genes downregulated) with an enrichment score >2. (M-N) Examples of genes involved in ECM organization (M) and Wnt signaling (N) that are only regulated in WT muscle. Data from 5 biological replicates (except for panels A-C). Data represents means ± SEM (if not otherwise stated). Statistics of RNAseq data were performed within the CLC genomics workbench software. Two-tailed Student t-test were performed in panel (A-C) to assess differences between untrained and trained animals as well as between genotypes. A paired two-tailed Student t-test was performed in (C) to observe individual improvements in VO_2max_. * indicates difference to Ctrl (pre-exercise condition) if not otherwise indicated; # indicates differences to the same condition in WTs; *<0.05; **<0.01; ***<0.001. See also Figure S5; Table S7.

Interestingly, some of the striking differences in acute gene expression between the untrained WT and mKO mice were mitigated by training. For example, the temporal difference, e.g. the shift towards peak gene expression at 0h, was also seen in the mKOs (Figure 5A). Additionally, the attenuation of the late response (4-8h post-exercise) is diminished in trained mKOs (Figures 5B-G). Moreover, less qualitative differences in transcript induction was seen in this context. Nevertheless, still 39% of the up-, and 62% of the downregulation was dependent on the presence of muscle PGC-1α in trained muscle (Figure 5H). These participate in encoding proteins involved in transcription, metabolism of lipids and carbohydrates, as well as ECM remodeling (Figures 5I, S6A; Table S7). Interestingly, the induction and repression of ECM genes in untrained and trained WT muscle following an acute endurance exercise bout, respectively, are under the control of PGC-1α (Figures 5I, S6B, S6C).

**Figure 5.**
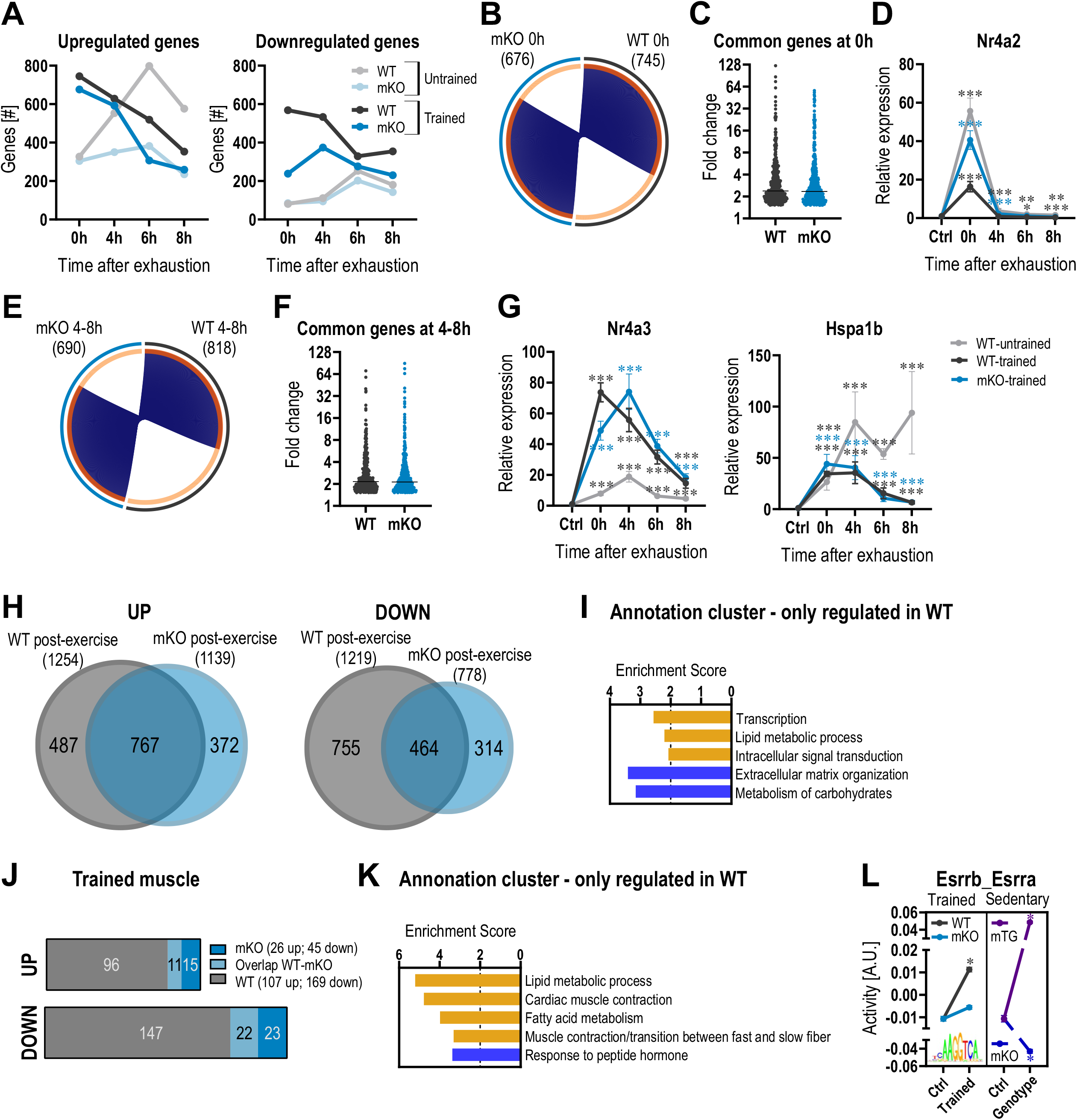
Differences in gene expression between WT and mKO animals is mitigated by training. (A) Number of genes that are up- and downregulated 0h, 4h, 6h and 8h after an acute exercise bout in untrained WT (light gray), trained WT (dark gray), untrained mKO (light blue) and trained mKO (dark blue) animals. (B) Circos plot of all upregulated genes immediately post-exercise (0h) in trained WT and mKO mice. (C) Relative fold change (and median) of all commonly regulated genes at the 0h time point in trained WT and mKO mice. (D) Example of gene trajectories with a peak immediately post-exercise in untrained WT, trained WT and trained mKO animals. (E) Circos plot of all upregulated genes after 4-8h (merged together) in trained WT and mKO mice. (F) Relative fold change (and median) of all commonly regulated genes 4-8h post-exercise in trained WT and mKO mice. (G) Examples of gene trajectories with a peak at a later time point in untrained WT, trained WT and trained mKO animals. (H) Venn diagrams of all up- and downregulated genes after an acute bout of exercise in trained WT (dark gray) and mKO (dark blue) mice. (I) All functional annotation clusters of up- (orange) and downregulated (blue) genes that are only regulated in trained WT mice (487 genes up- and 755 genes downregulated) with an enrichment score >2. (J) Bar Venn diagram of the genes altered in unperturbed trained WT (gray) and mKO (blue) muscle (overlap = light blue). (K) All functional annotation clusters of genes that are only up- (orange) and downregulated (blue) in trained muscle of WT animals only (up: n = 96; down = 147) with an enrichment score >2. (L) Motif of the transcription factors from ISMARA with the most significant activity change in trained WT animals and the comparison of the activity in trained mKO (left blue), gain-of-function model (sedentary muscle-specific PGC-1α transgenics – mTG – purple) and loss-of-function model (sedentary mKO, dark blue). Data from 5 biological replicates. Data represents means ± SEM (if not otherwise stated). Statistics of RNAseq data were performed within the CLC genomics workbench software. * indicates difference to Ctrl (pre-exercise condition); *<0.05 (for motif activity: * z-score >1.96); **<0.01; ***<0.001. See also Figure S6; Tables S4, S7.

Next, we investigated how these substantially altered transcriptional profiles after acute exercise bouts in untrained and trained muscles of PGC-1α propagate to the unperturbed trained quadriceps. First, the already constrained transcriptional changes in WT muscle were even further reduced in the absence of PGC-1α (Figure 5J). Of note, 90% of all up-, and 87% of all downregulated transcriptional events were dependent on the presence of PGC-1α in WT muscle (Figure 5J). Many of these genes encode proteins important for lipid metabolism and the fast-to-slow muscle fiber transition (Figure 5K; Table S7). As a coregulator, PGC-1α function depends on transcription factor binding to modulate gene expression. We therefore wanted to assess the mechanistic underpinnings of PGC-1α action in exercise. ISMARA revealed that in the context of acute exercise bouts, the absence of muscle PGC-1α affected 52% of all significantly affected motif activities in an untrained, and 39% in a trained muscle (Figures S6D-G; Table S4). Even more impressive, in the unperturbed trained muscle, almost all (91%) transcription factor binding motif activities were affected by the loss-of-function of PGC-1α in mKOs (Figure S6H; Table S4). Most notably, the significant training-linked increase in Esrrb_Esrra motif activity, a binding site for the estrogen-related receptor α (ERRα), was completed blunted in mKOs (Figure 5L). In line, the activity of this motif was highly increased in muscle-specific PGC-1α gain-of-function transgenic mice (mTG) and decreased in the loss-of-function model (Figure 5L). Thus, the massive constriction of transcription in the absence of muscle PGC-1α is associated with acute and persistent mitigations of transcription factor activities.

Finally, an abnormal endurance training adaptation was substantiated by the proteomic analysis of trained muscle of WT and mKO mice. First, many of the training-regulated proteins involved in mitochondrial respiration, lipid metabolic process and TCA cycle are already found at lower levels in sedentary mKO animals (Figures 6A, S7A; Tables S1, S2). In these mice, training fails to modulate such proteins as seen in WT controls (Figures 6B-E, S7B-D). For example, proteins involved in the TCA cycle, mitochondrial respiration, or reactive species detoxification were either not modulated by training, or did not reach the levels seen in sedentary WT muscle. The phenotypic, transcriptomic and proteomic data strongly indicate that PGC-1α thus is indispensable for a normal training response.

**Figure 6.**
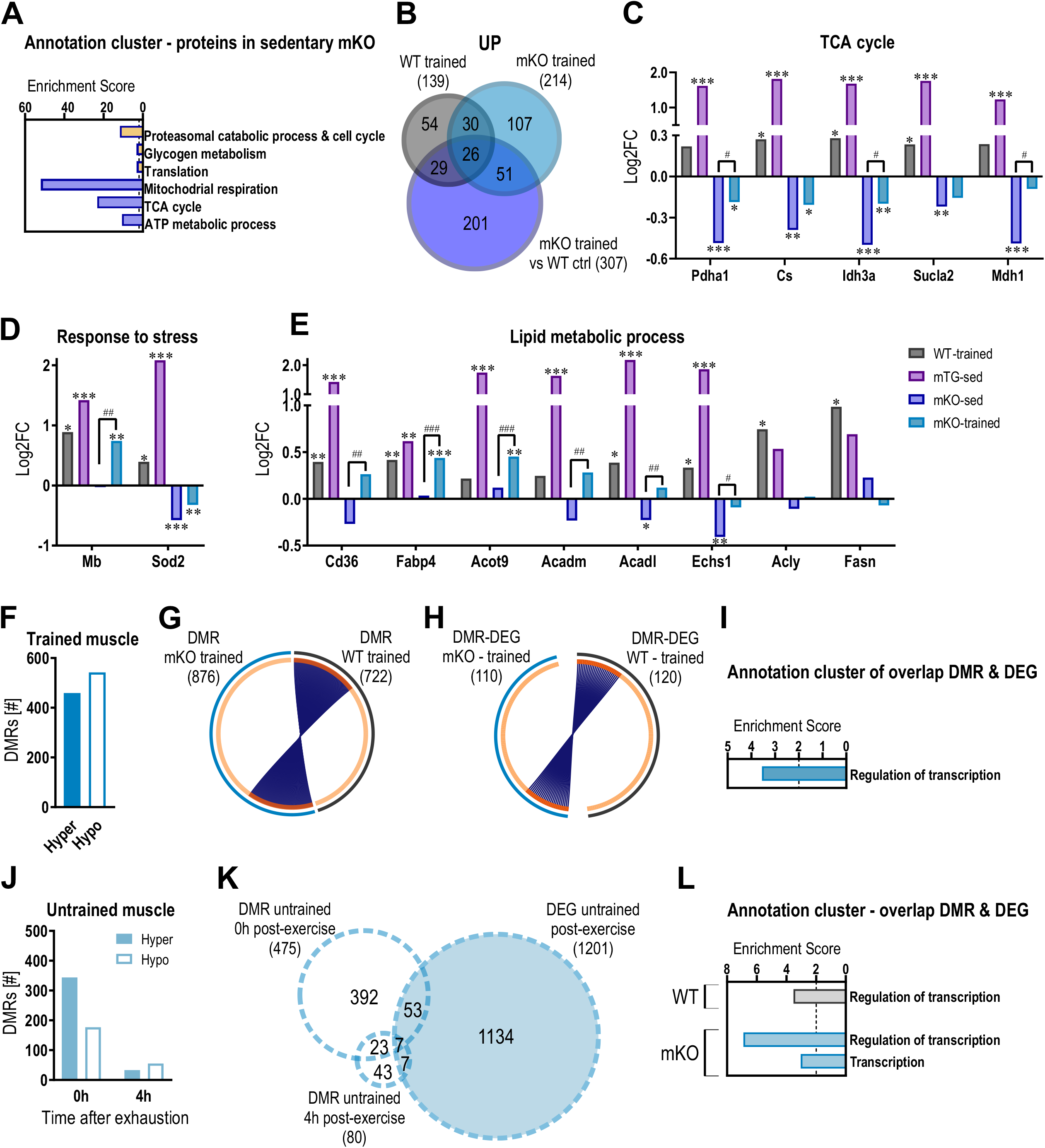
PGC-1α controls exercise-linked DNA methylation events. (A) Top 3 functional annotation clusters of up- (orange) and downregulated (blue) proteins in sedentary mKO compared to sedentary WT muscle. (B) Venn diagram of all upregulated proteins in trained WT (gray) and mKO muscle (lighter blue: trained mKO compared to sedentary mKO animals; darker blue: trained mKO compared to sedentary WT mice). (C-E) Examples of proteins involved in TCA cycle (C), response to stress (D) and lipid metabolic process (E) in trained WT (gray), sedentary mTG (pink), sedentary mKO (dark blue) and trained mKO (blue). Values are expressed relative to the WT sedentary control. (F) Number of differentially methylated regions (DMRs) in trained mKO muscle (hypermethylated = solid bar; hypomethylated = open bar). (G) Circos plot of all DMRs in trained mKO and those in trained WT animals. (H) Circos plot of differentially expressed genes (DEGs) with DMRs in trained WT and trained mKO animals (genes of the overlap from Figures 3H and S7E). (I) All functional annotation clusters of the 110 genes (overlap DMR and DEG from Figure S7E) in trained mKO animals with an enrichment score >2. (J) Number of hyper- (solid bars) and hypomethylated (open bars) regions 0h and 4h after exhaustion in untrained mKO animals. (K) Venn diagram of all DMRs 0h and 4h post-exercise (open circles) and DEGs after an acute bout of exercise (colored circle) in untrained mKO animals. (L) All functional annotation clusters of genes that are differentially methylated and transcriptionally regulated after an acute bout of exercise in WT (gray) and mKO (blue) mice. Data from 5-6 biological replicates. Statistics were performed using empirical Bayes moderated t-statistics for proteomics and within the CLC genomics workbench software for RNAseq data. * indicates difference to sedentary WT animals; # indicates difference to sedentary mKO animals; *<0.05; **<0.01; ***<0.001. See also Figure S7; Tables S1, S2, S6.

### PGC-1α controls exercise-linked DNA methylation events

We next compared the exercise- and training-induced epigenetic changes in WTs and mKOs. First, a markedly higher number of hypermethylated regions were found in trained mKO muscle, with little overlap with DMRs in WT quadriceps (Figures 6F, 6G). Similarly, the differentially expressed genes (DEGs) after acute exercise associated with DMRs of a trained muscle exhibited only a small overlap between the genotypes (Figures 6H, S7E). Nevertheless, many of these genes partitioned to regulation of transcription in mKOs, functionally similar to the results in WTs (Figures 6I, S7F, Table S6). Based on the largely different transcriptome of trained muscle, a divergence in DMRs might not be unexpected. However, it was surprising that absence of muscle PGC-1α also significantly altered epigenetic modulations in the untrained muscle after an acute exercise bout. At 0h, a relative shift from hypo-to hypermethylated regions was seen in mKO compared to WT mice (Figure 6J). DMRs almost completely disappeared at 4h (Figure 6J). Nevertheless, a strong functional cluster associated with transcription emerged from the overlap between DMRs and DEGs (Figures 6K, 6L; Table S6). Collectively, these results imply PGC-1α to be directly involved in the regulation of DNA methylation associated with gene expression. To test this hypothesis, we analyzed the epigenetic, transcriptomic and proteomic changes elicited by a muscle-specific PGC-1α gain-of-function model. Indeed, a substantial number of DMRs were detected in mTGs. Similar to trained WT quadriceps muscle, and mirroring the outcome in mKO animals, DMRs in mTGs skewed towards hypomethylation (Figure S7G). However, the overlap between DMRs of trained WT and sedentary mTG mice was very small and only 2.8% of the transcriptionally regulated genes have DMRs (Figures S7H, S7I). In line with previous observations^31^, the transcriptome of mTGs differs substantially from the chronically and acutely training- and exercise-regulated genes in WT muscle (Figure S7J). A better functional representation of training adaptation is provided by the mTG proteome, in which a strong accumulation of mitochondrial proteins, including members of the TCA cycle and respiratory chain, lipid metabolism, and a depletion of inflammation and proteasomal catabolic processes recapitulate many of the changes observed in trained WT muscle (Figure S7K; Table S1, S2).

## DISCUSSION

The ability of skeletal muscle to adapt to internal and external perturbations is a fundamental function, indispensable for human evolution as persistence hunters^32,33^. The plasticity evoked by endurance and resistance training leads to a pleiotropic remodeling of the function of many organs beyond muscle, linked to potent health benefits^1,6,7,34,35^. Inversely, the increasingly sedentary lifestyle in many societies defies the evolutionary changes, and thus constitutes a strong and independent risk factor for a large number of chronic pathologies, thereby contributing to morbidity and mortality^36,37^. While exercise-based interventions are highly efficacious in the prevention and therapy of many pathologies, often rivaling the effect of clinically approved drugs^6,38–40^, attempts to pharmacologically “mimic” training in a safe and efficient manner so-far have failed^41,42^. In light of the enormous fundamental and clinical significance of physical activity, it is surprising that our understanding of the underlying processes remains incomplete. Our findings now provide evidence for a much more complex process than proposed in prevailing models, with several novel aspects that describe muscle plasticity in different contexts as well as the basic mechanistic and regulatory principles of training adaptation.

First, even though massive morphological and functional remodeling is necessary for training adaptation, untrained and trained muscles are to a large extent transcriptionally indistinguishable, and steady-state gene expression changes explain only a small part of the corresponding modulation of the proteome. This was unexpected based on the contemporary view that repeated exercise bouts result in a steady increase in the basal expression of transcripts involved in mitochondrial function, substrate utilization, and other functional aspects that define a trained muscle^13^. Second, the massive, yet transient remodeling of the muscle transcriptome after acute exercise is quantitatively and qualitatively different when comparing a naïve to a trained muscle. Our findings vastly expand the prevailing models predicting an attenuation of the acute regulation of genes with repeated exercise bouts inasmuch we also report exacerbation, a shift in peak expression, and complete disappearance and *de novo* emergence of numerous transcripts (Figure 7). Finally, some transcripts exhibit diametrically opposite expression in acute exercise in untrained and trained muscles, e.g. genes encoding proteins involved in ECM remodeling, inflammation or axon guidance. This suggests a decidedly specific homeostatic perturbation and concomitant transcriptional response dependent on training status. These highly divergent modes of adaptation imply a complex regulatory framework by which training adaptation is brought about: expressed in the shift from a strong stress response and damage mitigation in naïve to improved resilience in trained muscle besides the metabolic, contractile and other adaptations. The deconvolution analysis however indicates that many of these changes are induced by events in non-muscle cells, in a presumably complex multicellular crosstalk and interaction. Future studies therefore have to consider this aspect and aim at an analysis at the level of individual cell types instead of bulk muscle tissue.

**Figure 7.**
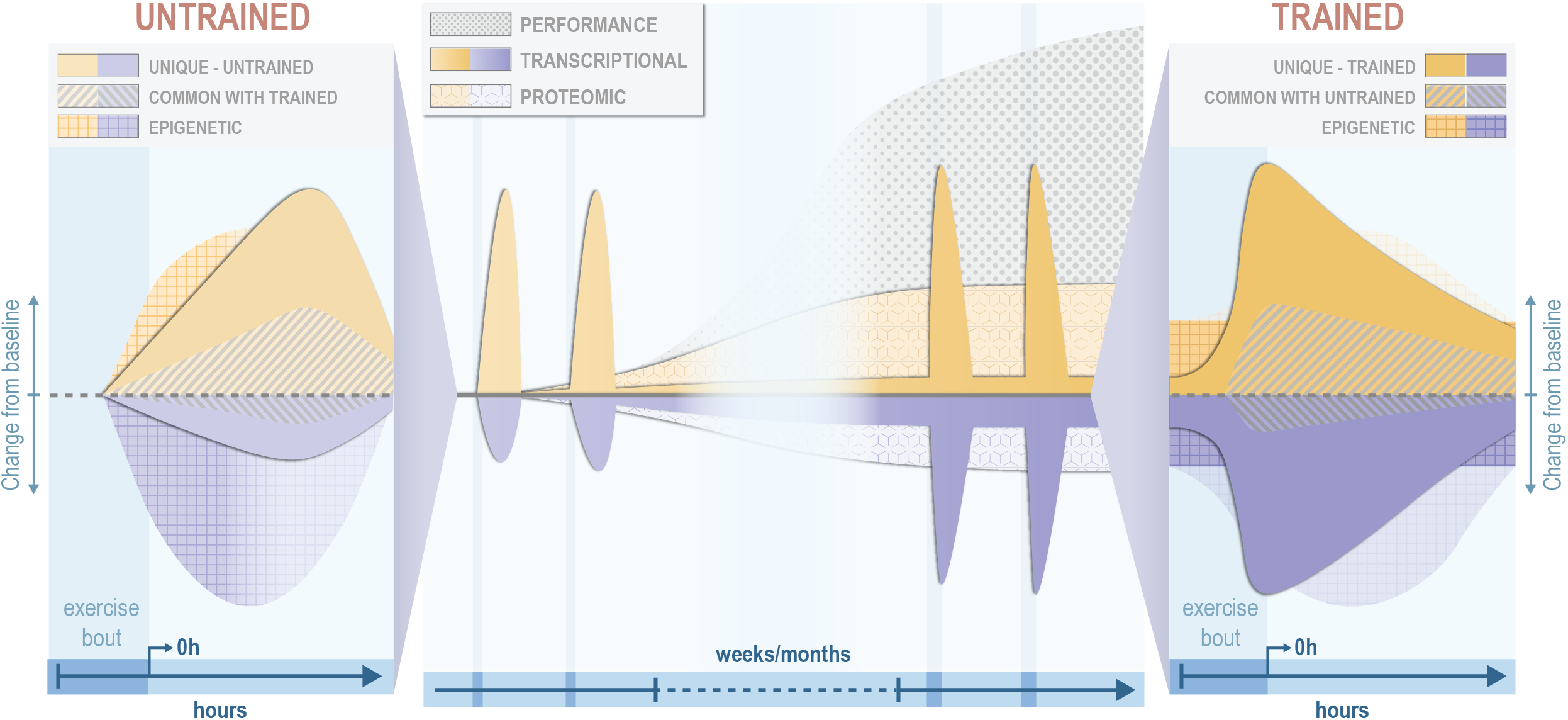
Schematic representation of the molecular exercise response. In an untrained muscle, short-term epigenetic regulation is coupled to acute perturbations of the transcriptome after an acute bout of exercise (orange = upregulated/hypermethylated; blue = downregulated/hypomethylated). With repeated bouts over time, a trained muscle is established hallmarked by morphological and functional adaptations that improve performance. This state is characterized by substantial proteomic remodeling, however in the context of a small number of chronically maintained gene expression modulation. Persistent modification of epigenetic marks prime the response of the trained muscle to recurring acute exercise bouts, in which substantial qualitative and quantitative changes in gene expression events compared to those observed in naïve muscle occur.

Our data also shed more light onto the mechanistic underpinnings of acute exercise and chronic training. We observed a clear differentiation between the acute epigenetic modifications and those persistently observed in a chronically trained muscle in an unperturbed state. The relatively small number of DMRs in close vicinity to differentially regulated genes in this context might be surprising, and might be at least in part caused by the limitation of RRBS. The association of epigenetic marks with gene expression, however, implies a priming of a limited number of key transcriptional regulators, which accordingly exhibit a different response to an acute bout of exercise in untrained and trained muscle. This priming might be sufficient for signal propagation and amplification to downstream genes and thereby contribute to the quantitative and qualitative differences in the transcriptional networks engaged in these two settings.

From the many regulatory factors that have been implied in exercise adaptation, we investigated the regulation and function of PGC-1α. We now unequivocally demonstrate that muscle PGC-1α is indispensable for normal transcriptional muscle plasticity, both after acute endurance exercise bouts in naïve and trained, as well as in the endurance-trained muscle. Moreover, we now show that VO_2max_, a marker for maximal endurance capacity, fails to significantly improve in the mKOs. Furthermore, training-induced shifts in the metabolism of ketone bodies and lactate are minimized in these animals^43,44^. Thus, collectively, these constraints might contribute to the limited gains that are possible in the absence of muscle PGC-1α, in our and previous studies resulting in a stagnation of endurance adaptation in mKOs at the level of untrained WT mice. Unexpectedly, we also found a strong impact of muscle PGC-1α on epigenetic marks, both chronically as well as acutely, both in loss-as well as in gain-of-function experiments. Future studies should therefore aim at investigating the molecular underpinnings of this link. Collectively, these findings demonstrate that regulatory factors such as PGC-1α, even though only acutely regulated, have a profound impact on longterm training adaptations. However, the regulatory complexity of muscle plasticity might have been underestimated since redundant, alternative or contingency pathways and factors seem to be able to be engaged in such settings to re-establish adaptation. This is not only true for PGC-1α, but also for AMPK and mTOR, which seem dispensable for certain aspects of training-induced muscle changes^45–48^. Such a complex regulatory framework would make sense in light of the evolutionary importance of the regulation of muscle plasticity, which has to function at least suboptimally to ensure survival even if individual factors fail.

Overall, our findings provide a new, refined, and much more complex model to describe how training adaptations are brought about. These results provide insights into an unsuspected and hitherto undescribed complexity in transcriptomic, epigenetic and proteomic changes in muscle plasticity, and hint at a vast, multifaceted mechanistic framework that controls the effects of acute exercise perturbations and long-term training alterations (Figure 7). Once validated and expanded in other species, muscles, training paradigms and time points, and in a more fine-grained cell type-specific manner, these insights will not only help to better understand such a fundamental process that was a main driver of human evolution, but also to leverage novel results to design strategies to benefit human health and well-being. It is encouraging that such efforts currently are ongoing, e.g. in the framework of the Wu Tsai Human Performance Alliance or the Molecular Transducers of Physical Activity Consortium (MoTrPAC)^49^.

## MATERIALS AND METHODS

A detailed description of the materials and methods are provided in the supplemental information.

## ACKNOWLEDGEMENTS

We thank Dekel Wainer and 4omix.com for the shiny app application, Prof. Markus Rüegg for the critical feedback on the manuscript and the Genomics Facility Basel, sciCORE, Proteomics Core Facility and the animal facility caretakers of the Biozentrum for their help. This work was supported by grants from the Swiss National Science Foundation, Switzerland (grant 310030_184832), the European Research Council Consolidator grant 616830-MUSCLE_NET, Siemens Fellowship for Excellence and the University of Basel.

## AUTHOR’S CONTRIBUTIONS

Conceptualization: RF, BH, JW and CH; Methodology: RF, BH, KJVN, DR, JW and CH; Investigation: RF, BH, SS, SAS; Analysis and interpretation: RF, BH, KJVN, SD, DR, VA and CH; Resources: JW and CH; Funding acquisition: RF and CH; Supervision: CH; Writing - original draft: RF and CH; Writing - reviews and editing: BH, KJVN, SS, SD, DR, VA, and JW.

## DATA AND CODE AVAILABILITY

Transcriptomic and RRBS data have been deposited at the Gene Expression Omnibus (GEO, accession numbers *XXX* and *YYY*, respectively). The transcriptomic data are furthermore accessible in an analyzed form at the myo-transcriptome of exercise database (myoTrEx, *LINK*). Proteomic data have been deposited at the Proteomics Identifications Database (PRIDE, accession number *ZZZ*). All other data are available upon reasonable request.

