## Supplemental Figures and Methods for "Molecular control of endurance training adaptation in mouse skeletal muscle"

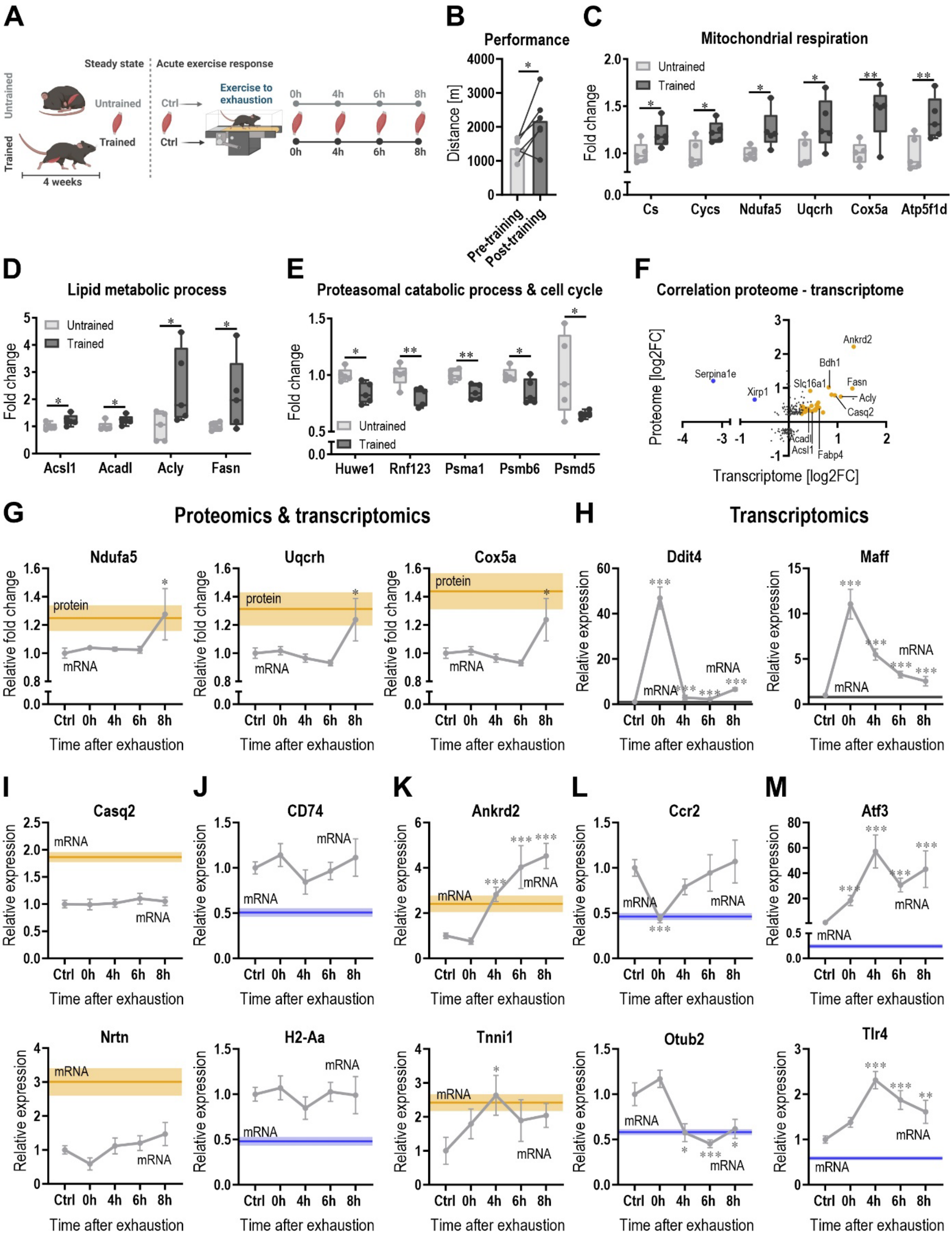


**Figure S1**

**Figure S1. The transcriptome of a trained is similar to that of an untrained muscle, related to Figure 1.**

(A) Schematic representation of the experimental setup.

(B) Changes in maximal running distance after 4 weeks of progressive treadmill training (n = 6 per group).

(C-E) Examples of proteins represented in the annotations clusters of mitochondrial respiration (C), lipid metabolic process (D) and proteasomal catabolic process (E) in a sedentary untrained muscle (light gray) and after 4 weeks of exercise training (dark gray).

(F) Correlation plot of the transcriptome and proteome of trained muscle. Only depicting genes/proteins that are significantly altered in the proteomics analysis of a trained muscle (cutoff: $p$ <0.05; Log2FC ±0.2). The colored (orange = up; blue = down) genes/proteins are also significantly altered on a transcriptional level in a trained muscle (FDR <0.05; Log2FC ±0.2).

(G) Representative proteins involved in aerobic respiration that are significantly increased in a trained muscle (solid orange line = mean ± SEM with light color) and acutely regulated on a transcriptional level post-exercise (gray line).

(H-M) Representative genes that are (H) only regulated after an acute exercise bout (light gray line; dark gray straight line represents the unchanged mRNA level in the trained muscle), (I & J) only changed in trained muscle (upregulated in (I) and downregulated (J) represented by the straight solid line in orange and blue, respectively), (K) upregulated in both trained muscle as well as after an acute bout of exercise (orange solid line represents the mean mRNA level in a trained muscle and the light gray line the changes after an acute bout of exercise), (L) downregulated in response to an acute bout of exercise and after training (blue line represents the mean mRNA level in a trained muscle and the light gray light the acute changes), and (M) upregulated after an acute challenge and downregulated after training (blue line represents the downregulation in a trained muscle and the light gray line the acute changes). The light gray line represents the acute changes after one exercise bout and the straight solid line represents the mean values in the unperturbed trained muscle (± SEM in lighter color); line color of trained muscle: orange = significantly upregulated; blue = significantly downregulated; gray = unchanged in unperturbed trained muscle.

Data from 5 biological replicates (if not otherwise stated). Data represents means ± SEM. Statistics were performed using empirical Bayes moderated t-statistics for proteomics and within the CLC genomics workbench software for RNAseq data. For the running distance (B) a paired two-tailed Student's t-test was performed. * indicates difference to Ctrl (pre-exercise condition) if not otherwise indicated; *<0.05; **<0.01; ***<0.001. See also Tables S1, S2.

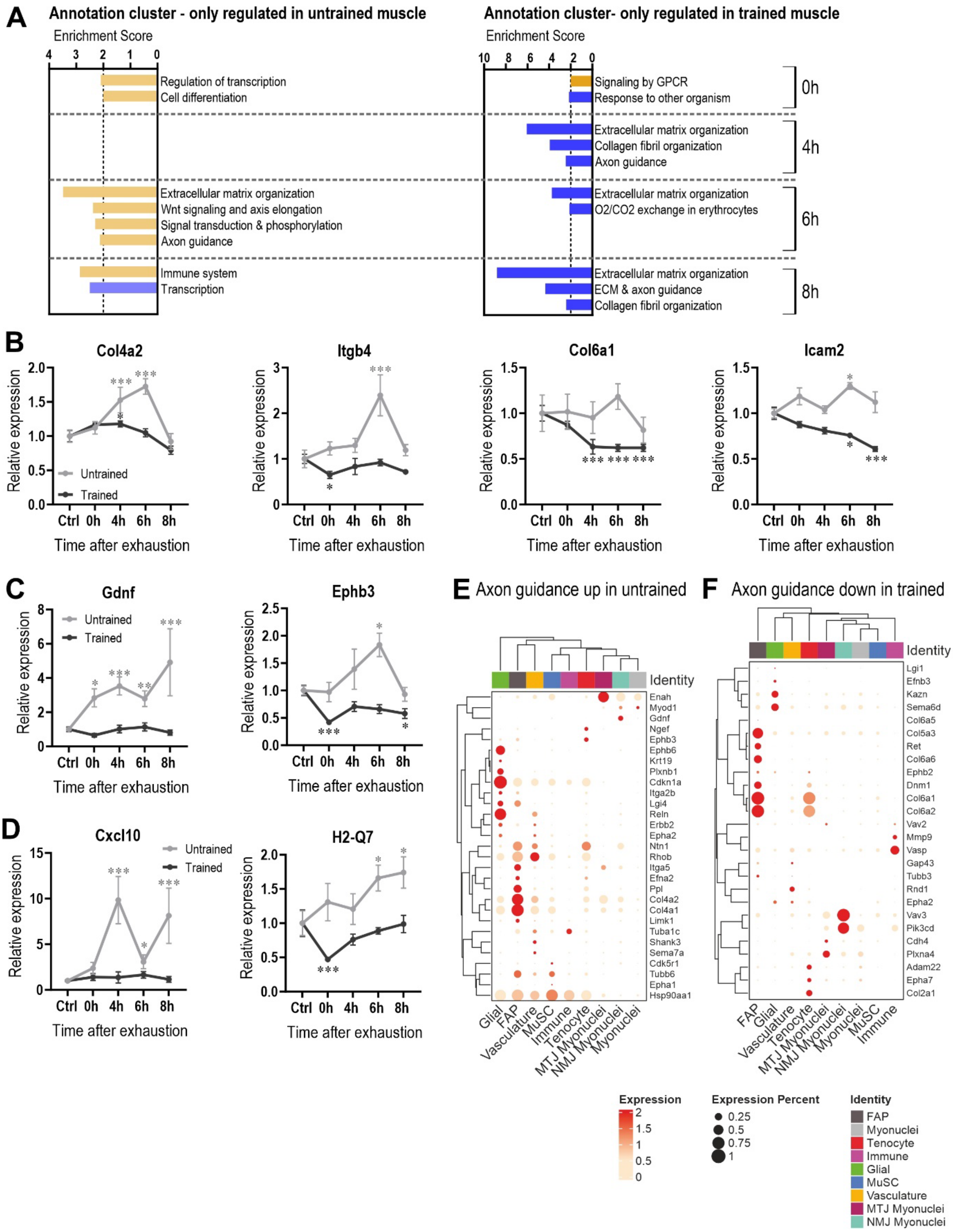
A
Annotation cluster - only regulated in untrained muscle
Enrichment Score
Regulation of transcription
Cell differentiation
Extracellular matrix organization
Wnt signaling and axis elongation
Signal transduction & phosphorylation
Axon guidance
Immune system
Transcription
Annotation cluster- only regulated in trained muscle
Enrichment Score
Signaling by GPCR
Response to other organism
Extracellular matrix organization
Collagen fibril organization
Axon guidance
Extracellular matrix organization
O2/CO2 exchange in erythrocytes
Extracellular matrix organization
ECM & axon guidance
Collagen fibril organization
0h
4h
6h
8h
B
Col4a2
Itgb4
Col6a1
Icam2
Relative expression
Untrained
Trained
Ctrl 0h 4h 6h 8h
Time after exhaustion
C
Gdnf
Ephb3
D
Cxcl10
H2-Q7
E Axon guidance up in untrained
F Axon guidance down in trained
Identity
Expression
Expression Percent
FAP
Myonuclei
Tenocyte
Immune
Glial
MuSC
Vasculature
MTJ Myonuclei
NMJ Myonuclei

**Figure S2**

**Figure S2. Qualitative transcriptional response to exercise depends on training status, related to Figure 2.**

(A) All functional annotation clusters of the up- (orange) and downregulated (blue) genes with an enrichment score >2 that are only regulated the untrained (left; light colors) or trained (right; darker colors) muscle after an acute exercise bout per time point.

(B-D) Examples of gene trajectories in untrained (light gray) and trained (dark gray) muscle involved in ECM organization (B), axon guidance (C) and immune system (D).

(E-F) Deconvolution of genes involved in axon guidance that are upregulated in an untrained (E) and downregulated in a trained (F) muscle.

Data from 5 biological replicates. Data represents means ± SEM. Statistics of RNAseq data were performed within the CLC genomics workbench software. * indicates difference to Ctrl (pre-exercise condition); *<0.05; **<0.01; ***<0.001. See also Table S3.

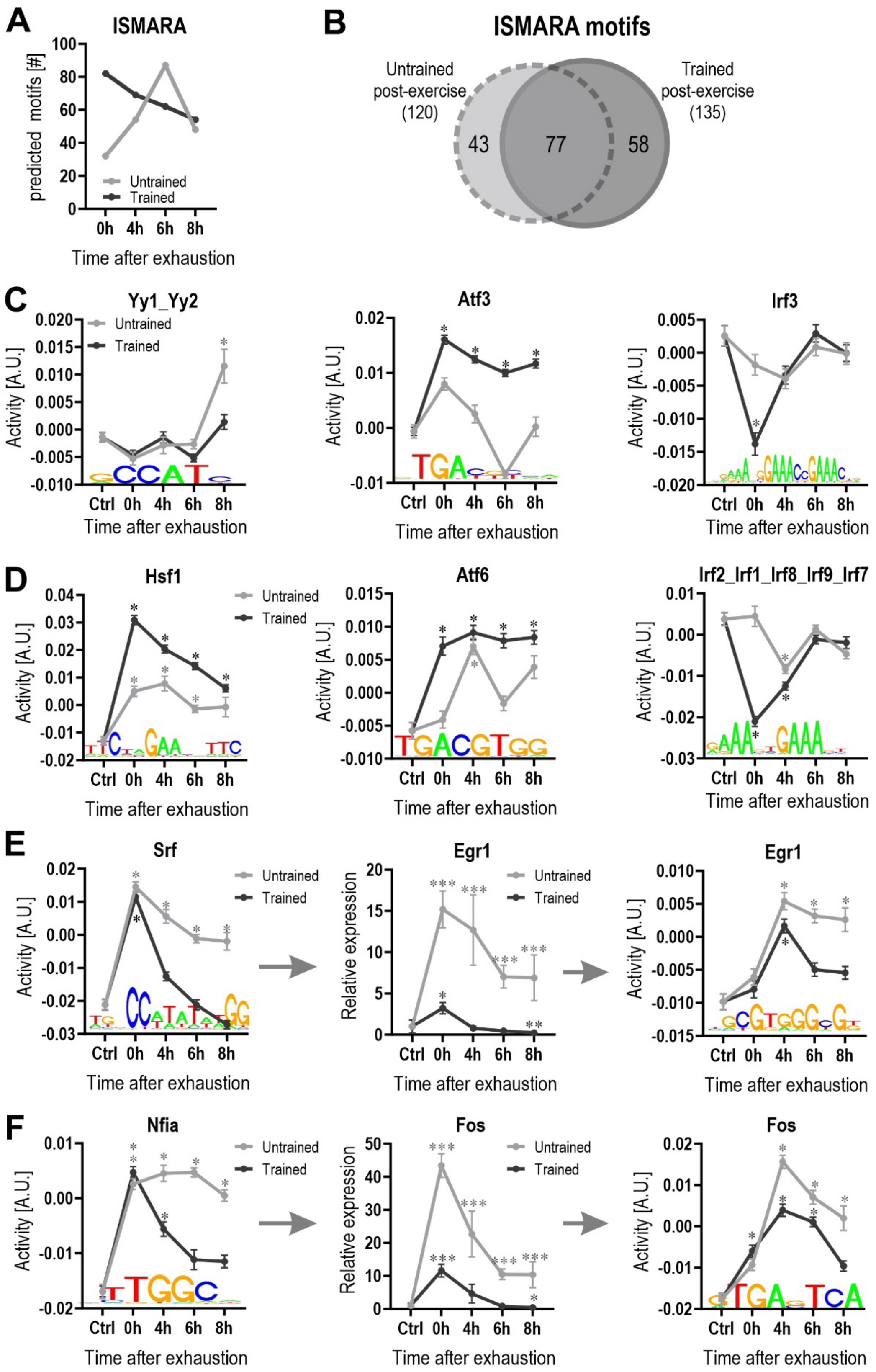
A
ISMARA
predicted motifs [#]
Untrained
Trained
Time after exhaustion
B
ISMARA motifs
Untrained post-exercise (120)
Trained post-exercise (135)
43
77
58
C
Yy1_Yy2
Atf3
Irf3
Activity [A.U.]
D
Hsf1
Atf6
Irf2_Irf1_Irf8_Irf9_Irf7
E
Srf
Egr1
Relative expression
F
Nfia
Fos

**Figure S3**

**Figure S3. Faster transcriptional response in trained muscle after one bout of exhaustion exercise, related to Figure 3.**

(A. Number of motifs of transcription factors from ISMARA that have an increased or decreased activity (z-score >1.96) per time point in an untrained (light gray) and trained (dark gray) muscle after an acute exercise bout.

(B) Venn diagram of all the predicted motifs that are changes in untrained (light gray, dashed line) and trained (dark gray, solid line) muscle after an acute exercise bout.

(C-D) Trajectories of motif activity of transcription factors from ISMARA in untrained (light gray) and trained (dark gray) muscle post-exercise that are either specific to training status (C) or show an exacerbation after training (D).

(E-F) Examples of possible transcriptional cascades including top predicted transcription factors by ISMARA and their downstream targets (expression and motif activities).

Data from 5 biological replicates. Data represents means ± SEM. Statistics of RNAseq data were performed within the CLC genomics workbench software. * indicates difference to Ctrl (pre-exercise condition); *<0.05 (for motif activity: * z-score >1.96); **<0.01; ***<0.001. See also Table S4.

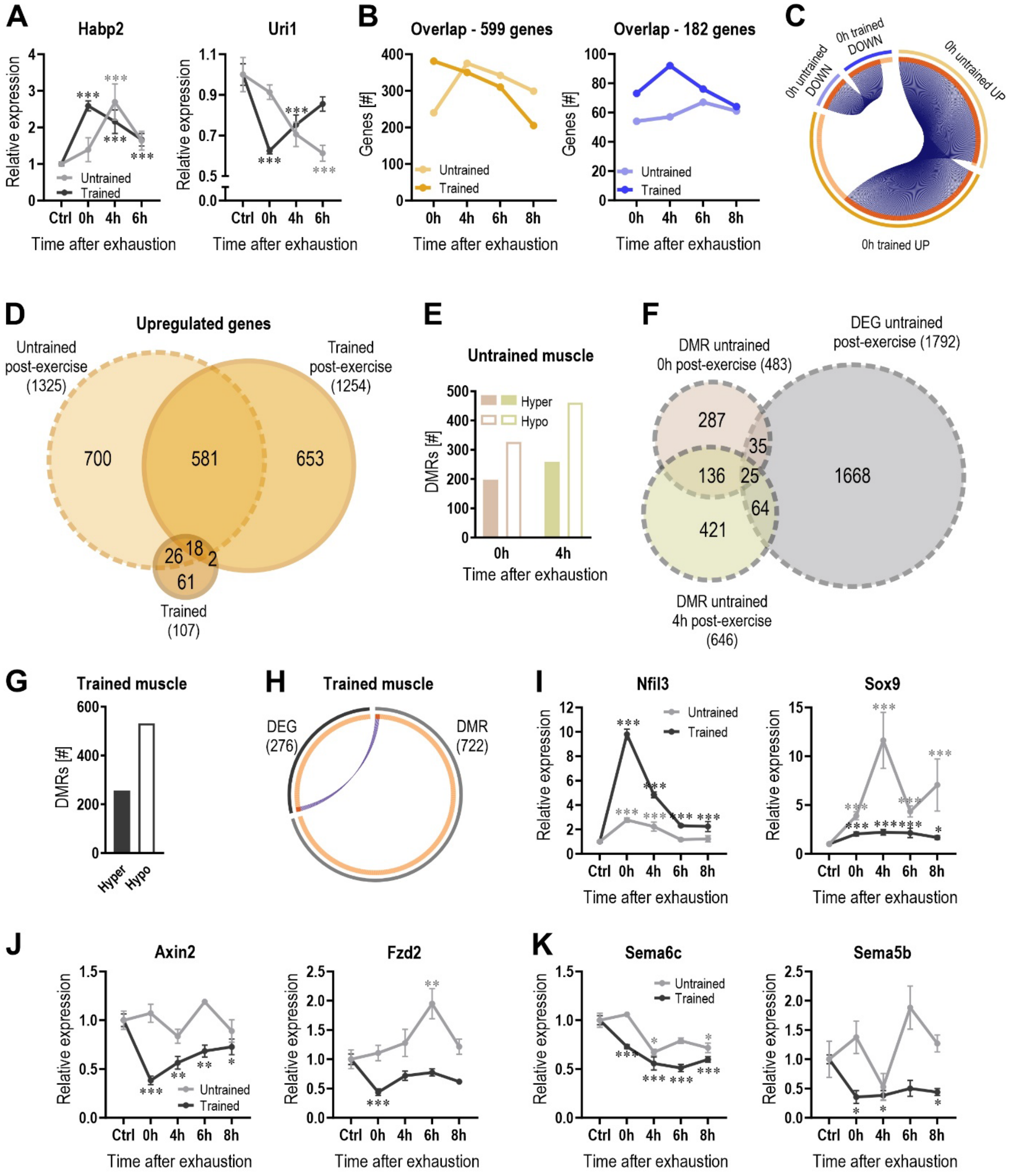
A
Habp2
Uri1
Relative expression
Untrained
Trained
Ctrl 0h 4h 6h
Time after exhaustion
B
Overlap - 599 genes
Overlap - 182 genes
Genes [#]
0h 4h 6h 8h
C
0h untrained DOWN
0h trained DOWN
0h untrained UP
0h trained UP
D
Upregulated genes
Untrained post-exercise (1325)
Trained post-exercise (1254)
Trained (107)
700
581
653
26
18
2
61
E
Untrained muscle
Hyper
Hypo
DMRs [#]
F
DMR untrained 0h post-exercise (483)
DEG untrained post-exercise (1792)
DMR untrained 4h post-exercise (646)
287
35
136
25
1668
64
421
G
Trained muscle
Hyper
Hypo
H
Trained muscle
DEG (276)
DMR (722)
I
Nfil3
Sox9
J
Axin2
Fzd2
K
Sema6c
Sema5b
Ctrl 0h 4h 6h 8h

**Figure S4**

**Figure S4. Faster transcriptional response in trained muscle after one bout of exhaustion exercise, related to Figure 3**.

(A) Representative examples of genes with the same maximal/minimal expression change but with an accelerated induction/repression in gene expression (phase shift) in trained muscle after acute exercise (dark gray) compared to untrained (light gray).

(B)Visualization of the temporal trajectories of the overlapping genes from Figure 2B in untrained (light color) and trained (dark color) muscle (orange = upregulated; blue = downregulated).

(C) Circos plot of the genes at the 0h time point of (B) to visualize the large overlap between the genes that are regulated immediately post-exercise.

(D) Venn diagram of all upregulated genes after an acute exercise bout in untrained (light orange, dashed line) and trained (orange, solid line) muscle as well as in an unperturbed trained muscle (dark orange).

(E) Number of hyper- (solid bars) and hypomethylated (open bars) regions 0h and 4h after exhaustion in an untrained muscle.

(F) Venn diagram of all differentially methylated regions (DMRs) 0h and 4h post-exercise (colored circles) and the differentially regulated genes (DEGs) after an acute bout of exercise (gray circle).

(G) Number of DMRs in an unperturbed trained muscle (hypermethylated = solid bar; hypomethylated = open bar).

(H) Circos plot of all DMRs and DEG in an unperturbed trained muscle.

(I-K) Examples of differentially methylated genes involved in transcription (I), Wnt signaling (J) and axon guidance (K) in untrained (light gray) and trained (dark gray) muscle post-exercise.

Data from 5 biological replicates. Data represents means ± SEM. Statistics of RNAseq data were performed within the CLC genomics workbench software. * indicates difference to Ctrl (pre-exercise condition); *<0.05; **<0.01; ***<0.001. See also Table S6.

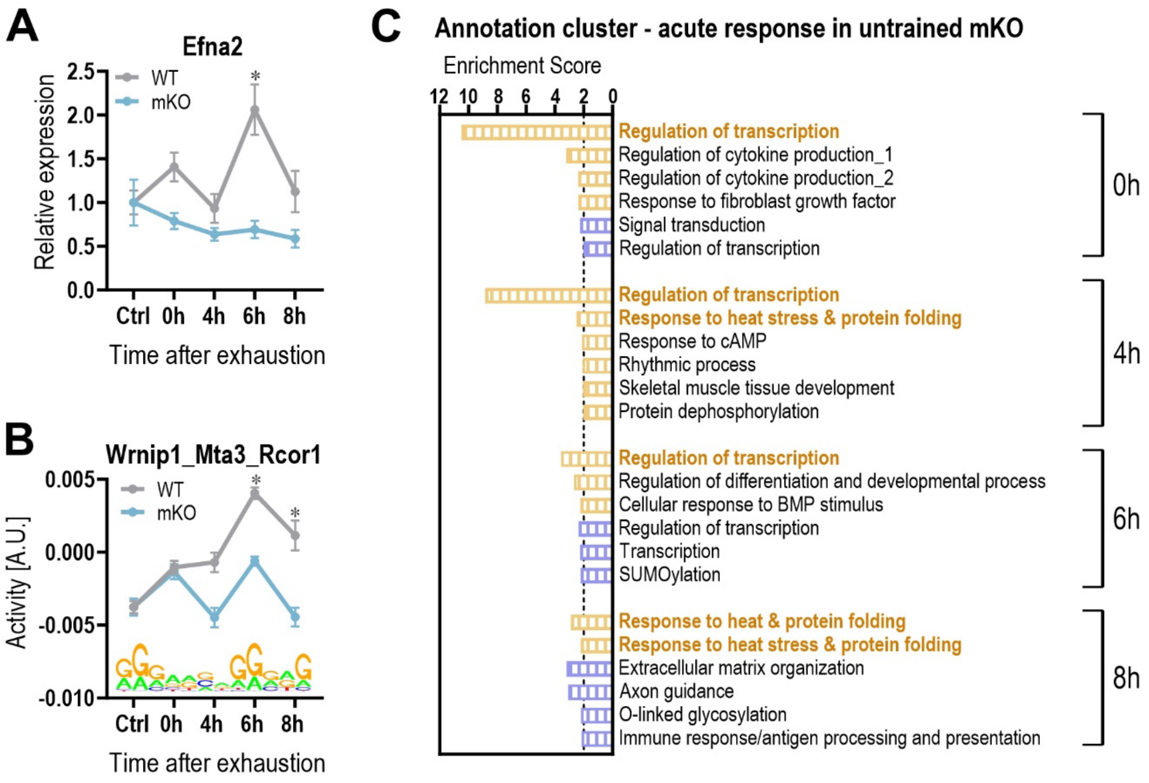


**Figure S5**

**Figure S5. PGC-1α is indispensable for a normal transcriptional response to acute exercise and long-term training, related to Figure 4.**

(A) Example of transcriptional trajectories of a gene involved in microglial cell proliferation in untrained WT (light gray) and mKO (light blue) muscle post-exercise.

(B) Prediction of the activity of a motif using ISMARA that might be involved in the regulation of ECM-related genes.

(C) All annotation clusters of genes that are significantly up- (orange) or downregulated (blue) in mKO animals after an exercise bout with an enrichment score >2.

Data from 5 biological replicates. Data represents means ± SEM. Statistics of RNAseq data were performed within the CLC genomics workbench software. * indicates difference to Ctrl (pre-exercise condition); *<0.05 (for motif activity: * z-score >1.96); **<0.01; ***<0.001. See also Tables S4, S7.

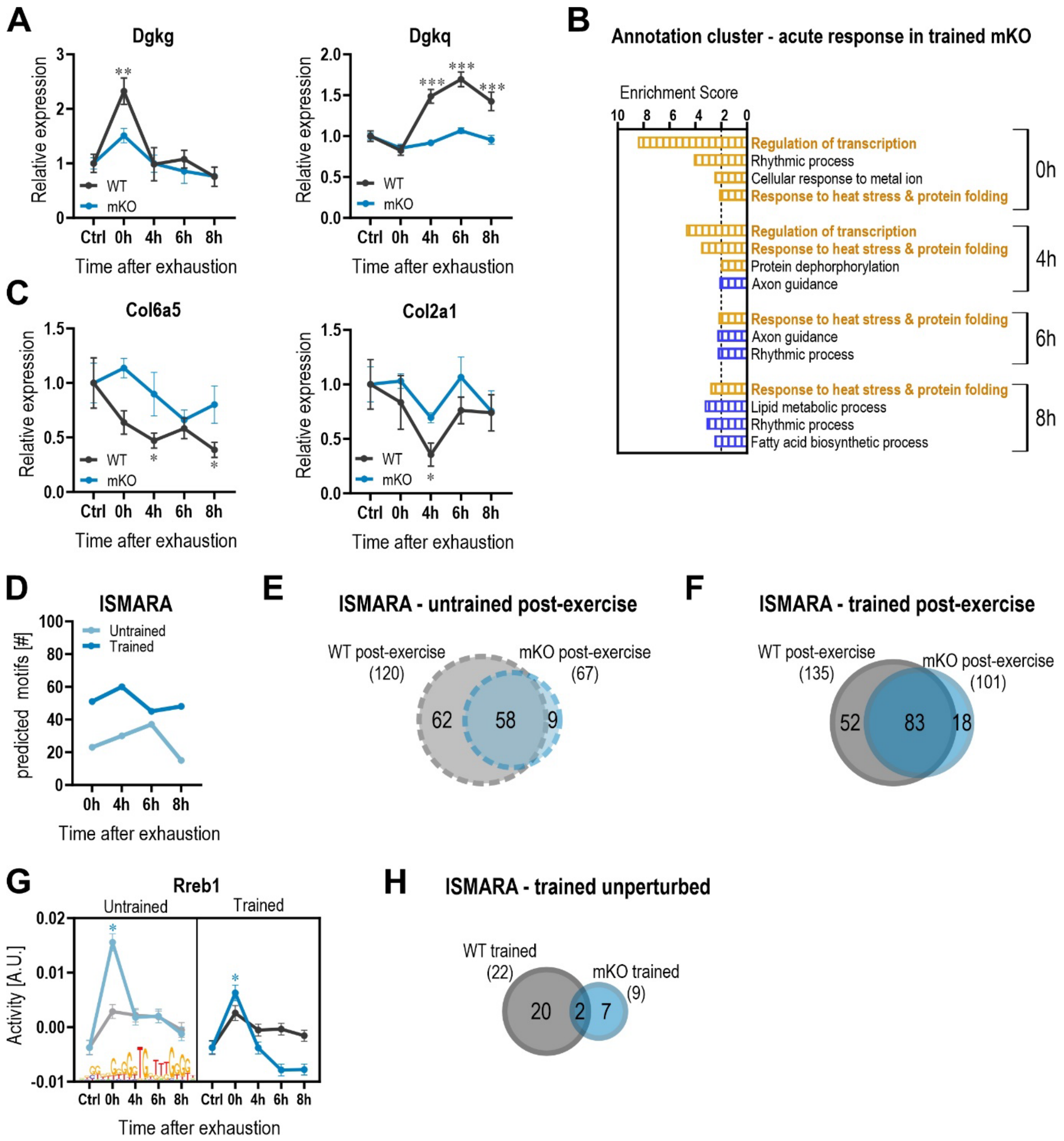
A
Dgkg
Relative expression
WT
mKO
Ctrl 0h 4h 6h 8h
Time after exhaustion
Dgkq
B
Annotation cluster - acute response in trained mKO
Enrichment Score
Regulation of transcription
Rhythmic process
Cellular response to metal ion
Response to heat stress & protein folding
0h
Regulation of transcription
Response to heat stress & protein folding
Protein dephorphorylation
Axon guidance
4h
Response to heat stress & protein folding
Axon guidance
Rhythmic process
6h
Response to heat stress & protein folding
Lipid metabolic process
Rhythmic process
Fatty acid biosynthetic process
8h
C
Col6a5
Col2a1
D
ISMARA
Untrained
Trained
predicted motifs [#]
E
ISMARA - untrained post-exercise
WT post-exercise (120)
mKO post-exercise (67)
62
58
9
F
ISMARA - trained post-exercise
WT post-exercise (135)
mKO post-exercise (101)
52
83
18
G
Rreb1
Untrained
Trained
Activity [A.U.]
H
ISMARA - trained unperturbed
WT trained (22)
mKO trained (9)
20
2
7

**Figure S6**

**Figure S6. Differences in gene expression between WT and mKO animals is mitigated by training, related to Figure 5**.

(A) Examples of genes involved in lipid metabolic process in trained WT (gray) and mKO (blue) mice.

(B) All annotation clusters of genes that are significantly up- (orange) or downregulated (blue) in trained mKO animals after an exercise bout with an enrichment score >2.

(C) Examples of genes involved in ECM organization in trained WT (gray) and mKO (blue) mice.

(D) Number of motifs of transcription factors from ISMARA that have an increased or decreased activity (z-score >1.96) per time point in an untrained (light) and trained (dark) mKO muscle after an acute exercise bout.

(E-F) Venn diagram of all predicted motifs that are changes after an acute exercise bout in untrained (E) and trained (F) WT (gray) and mKO (blue) animals.

(G) Trajectories of the activity of a motif that is only altered in the absence of muscle PGC-1α.

(H) Venn diagram of all predicted motifs that are changes in unperturbed trained WT (gray) and mKO (blue) muscle.

Data from 5 biological replicates. Data represents means ± SEM. Statistics of RNAseq data were performed within the CLC genomics workbench software. * indicates difference to Ctrl (pre-exercise condition); *<0.05 (for motif activity: * z-score >1.96); **<0.01; ***<0.001. See also Tables S4, S7.

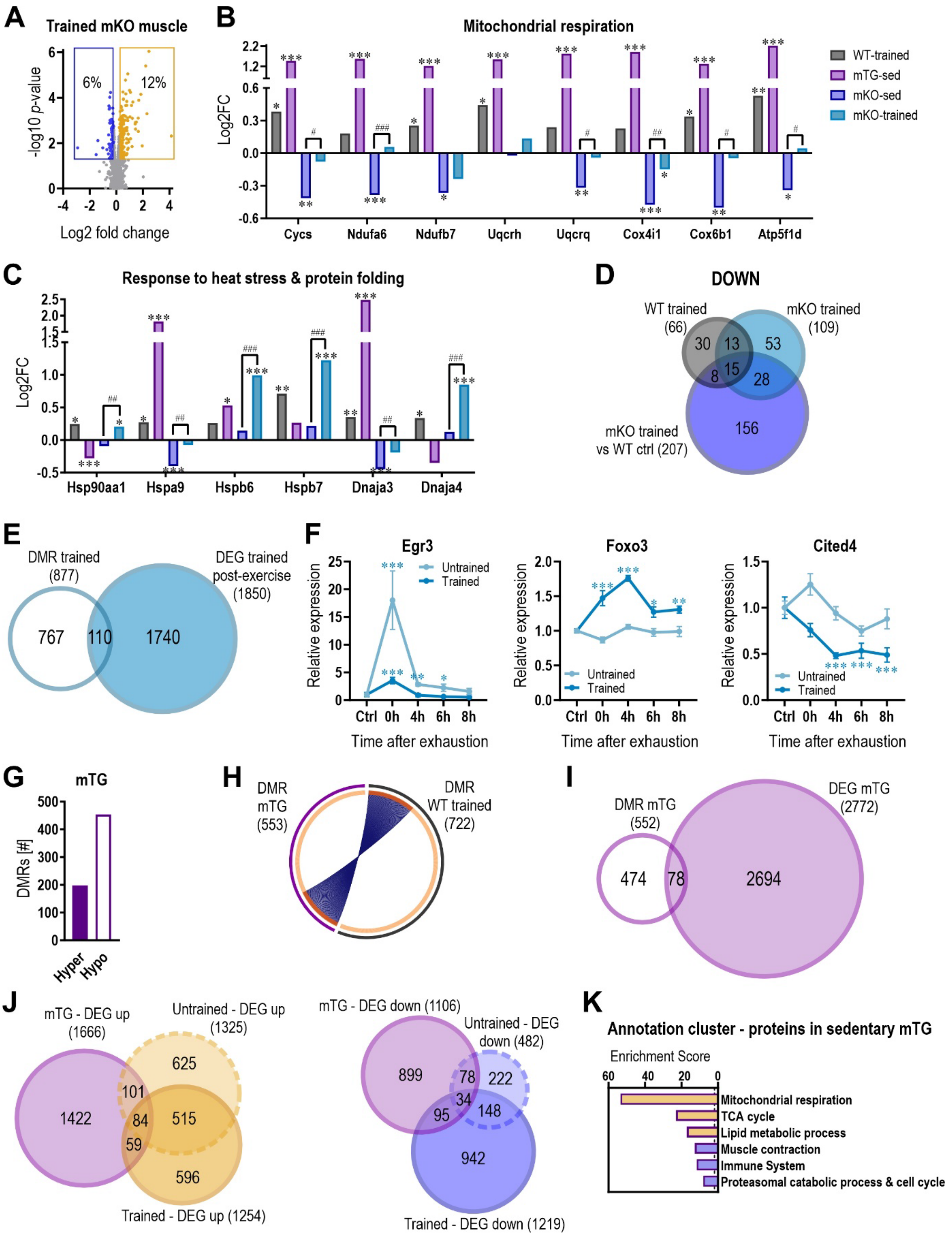
A
Trained mKO muscle
6%
12%
-log10 p-value
Log2 fold change
B
Mitochondrial respiration
Log2FC
Cycs
Ndufa6
Ndufb7
Uqcrh
Uqcrq
Cox4i1
Cox6b1
Atp5f1d
WT-trained
mTG-sed
mKO-sed
mKO-trained
C
Response to heat stress & protein folding
Log2FC
Hsp90aa1
Hspa9
Hspb6
Hspb7
Dnaja3
Dnaja4
D
DOWN
WT trained (66)
mKO trained (109)
mKO trained vs WT ctrl (207)
30
13
53
8
15
28
156
E
DMR trained (877)
DEG trained post-exercise (1850)
767
110
1740
F
Egr3
Foxo3
Cited4
Untrained
Trained
Relative expression
Ctrl 0h 4h 6h 8h
Time after exhaustion
G
mTG
DMRs [#]
Hyper
Hypo
H
DMR mTG (553)
DMR WT trained (722)
I
DMR mTG (552)
DEG mTG (2772)
474
78
2694
J
mTG - DEG up (1666)
Untrained - DEG up (1325)
Trained - DEG up (1254)
1422
625
101
84
515
59
596
mTG - DEG down (1106)
Untrained - DEG down (482)
Trained - DEG down (1219)
899
78
222
34
95
148
942
K
Annotation cluster - proteins in sedentary mTG
Enrichment Score
Mitochondrial respiration
TCA cycle
Lipid metabolic process
Muscle contraction
Immune System
Proteasomal catabolic process & cell cycle

**Figure S7**

**Figure S7. PGC-1α controls exercise-linked DNA methylation events, related to Figure 6.**

(A) Volcano plot of all detected proteins in trained quadriceps muscle of mKO using mass spectrometry-based proteomics. Proteins that are significantly upregulated (cutoff: $p$ <0.05; Log2FC ±0.2) are depicted in orange while those that are lower abundant are shown in blue (relative to sedentary mKO mice).

(B-C) Examples of proteins involved in mitochondrial respiration (B) and in response to heat stress & protein folding (C) in trained WT (gray), sedentary mTG (pink), sedentary mKO (dark blue) and traind mKO (blue). Values are expressed relative to the WT sedentary control.

(D) Venn diagram of the proteins that are downregulated in a trained WT (gray) and mKO muscle (ligher blue: trained mKO compared to sedentary mKO animals; darker blue: trained mKO compared to sedentary WT mice).

(E) Venn diagram of all genes with differentially methylated regions (DMRs) in unperturbed trained mKO muscle and differentially expressed genes (DEGs) in trained mKO muscle upon an acute bout of exercise.

(F) Trajectories of transcription factors in untrained (light blue) and trained (blue) mKO mice that are differentially methylated after training.

(G) Number of DMRs in sedentary mTG muscle (hypermethylated = solid bar; hypomethylated = open bar).

(H) Circos plot of all DMRs in sedentary mTG and those in trained WT animals.

(I) Venn diagram of all genes with DMRs and those that are differentially expressed (DEG) in sedentary mTG muscle (compared to sedentary WT mice).

(J) Venn diagrams of all up- and downregulated genes in sedentary mTG muscle (pink) and after an acute bout of exercise in untrained (light orange or blue) and trained (darker orange and blue) muscle.

(K) Top 3 functional annotation clusters of the up- (orange) and downregulated (blue) proteins in sedentary mTG muscle compared to sedentary WT.

Data from 5-6 biological replicates. Data represents means ± SEM. Statistics were performed using empirical Bayes moderated t-statistics for proteomics and within the CLC genomics workbench software for RNAseq data. * indicates difference to sedentary WT animals; # indicates difference to sedentary mKO animals; *<0.05; **<0.01; ***<0.001. See also Tables S1, S2, S6, S7.

### SUPPLEMENTAL INFORMATION

### MATERIALS AND METHODS

#### *Animals*

For this study, C57BL/6 male mice lacking PGC-1α specifically in muscle (mKO) or transgenically overexpressing PGC-1α in muscle (mTG) were used. PGC-1α mKO mice were generated by breeding PGC-1α$^{flox/flox}$ mice with a HSA-cre mouse line (Jackson Laboratories stock number: 009666) as described previously[1,2]. For the generation of the PGC-1α mTG animals, C57BL/6 mice expressing PGC-1α under the control of the creatine kinase promoter were crossed with WT mice as described previously[3]. Their respective littermates served as WT control animals. Mice had free access to water and a standard rodent chow diet and were housed under standard conditions with a 12h light/12h dark cycle. The experiments were performed with 5-6 mice per condition and mice were sacrificed at the age of 18-24 weeks. All experimental protocols followed the Swiss guidelines for animal experimentation and care and were approved by the Kantonales Veterinäramt Basel-Stadt.

#### *Exercise protocols*

Exercise training was performed on a motorized treadmill (Columbus Instruments) on 5 days per week for 1h. The training protocol was progressive with increasing velocity from 10 m/min to 20 m/min at an inclination of 5°. Trained mice were sacrificed either 18h after the last training session and used as steady state condition of a trained muscle or they performed a maximal performance test after a 72h rest period. As a control for the acute exercise response of trained mice, a group of trained mice was sacrificed 72h after the last training session, corresponding to the time point of the final maximal exercise bout.

Maximal performance and the acute exercise response to one bout of exhaustion exercise was assessed on a motorized treadmill (Columbus Instruments) as described previously[4]. Prior to the test, mice were familiarized with treadmill running for two days. The maximal exercise test was performed at an inclination of 5° and after a warming up of 5 min at 5 m/min followed by 5 min at 8 m/min, the velocity was progressively increased 2 m/min every 15 min until exhaustion. To determine the acute exercise response, mice were sacrifice and tissue collected either immediately (0h) post-exercise or after 4h, 6h or 8h (Figure S1A). The collected tissue was snap frozen in liquid nitrogen and stored at -80°C for further analyses.

Maximal oxygen consumption ($VO_{2max}$) was measured during a short maximal exercise test on a closed treadmill (Columbus Instruments) with only a subset of mice. Similar to the maximal performance test, mice were first familiarized with treadmill running for two days. The test was performed at an inclination of 15°. After 5 min of acclimatization to the closed treadmill chamber at 0 m/min, the velocity was increased to 10 m/min and then increased every 2 min with 2 m/min until exhaustion. After the test, mice were put back into their home cage.

*RNA sequencing (RNAseq) and data analysis*

After homogenizing pulverized quadriceps muscle in 1 ml of TRIzol agent (Sigma-Aldrich) using FastPrep tubes (MP Biomedicals), RNA was isolated according to the manufacturer's protocol. RNA concentration and quality were measured on the NanoDrop OneC spectrophotometer (Thermo Fisher Scientific). Subsequently, 7'500 ng RNA was further purified using the Direct-zol RNA MiniPrep Kit (Zymo Research). Following the RNAseq library preparation with 1 µg of purified RNA using the TruSeq RNA library Prep Kit (Illumina) according to the manufacturer's instructions, single read sequencing was performed on the HighSeq 2500 machine (50 cycles, Illumina).

Next, fastq files were mapped to the mouse genome (mm10) and differential gene expression analysis was performed using the CLC Genomics Workbench Software (version 21.0.5, Qiagen). For all downstream analysis, a Log2FC cutoff of ±0.6 (genes with a Log2FC >0.599 or <-0.599) and FDR value <0.05 was used. The prediction of enriched transcription factor binding motifs was done by Integrated Motif Activity Response Analysis (ISMARA; https://ismara.unibas.ch/mara/)[5]. The Database for Annotation, Visualization and Integrated Discovery (DAVID; https://david.ncifcrf.gov/tools.jsp) platform was used to determine functional annotation clusters of GO biological processes and REACTOME pathways and clusters with an enrichment score >2 were considered[6,7]. The overlap of genes were determined using InteractiVenn (http://www.interactivenn.net/index.html)[8] and results were visualized with Heatmaps using Morpheus (https://clue.io/morpheus), Circos plots using Metascape (https://metascape.org/gp/index.html#/main/step1)[9] or proportional Venn diagrams using DeepVenn (https://www.deepvenn.com/)[10].

*Genomic DNA isolation*

Approximately 15 mg of pulverized quadriceps muscle was used for genomic DNA (gDNA) isolation. Tissue was digested overnight in proteinase K (20 mg/ml) (Promega) and DNA lysis buffer (50 mM Tris-HCl pH-8.0, 100 mM NaCl, 10 mM EDTA, 0.5% Nonidet P-40) at 55°C on a shaker. The next day proteinase K was inactivated at 95°C for 10 min. Subsequently, phenol-chloroform-isoamyl alcohol (PCI) (Sigma-Aldrich) was added in a 1:1 ratio and the samples were vortexed and centrifuged at room temperature (RT) at 13’000 rpm for 4 min. Next, the upper phase was collected, the same volume of PCI as in the first step added, vortexed and centrifuged as described above. Then, the upper phase was collected again and 1/10 volume of 3M Na-Acetate pH 5.0 and 6/10 volume of Isopropanol was added. The samples were vortexed, incubated at RT for 5 min and centrifuged at RT at maximum speed (20’000 rpm) for 15 min. Subsequently, the supernatant was removed and the pellet washed with 70% ethanol and centrifuge at RT at maximum speed for 5 min. After removing the supernatant, the pellet was dried for 10 min at RT and resuspended in nuclease free $H_2O$. The gDNA quality and concentration were measured on the NanoDrop OneC spectrophotometer (Thermo Fisher Scientific). The isolated gDNA was further purified according to the manufacturer’s instructions using the DNeasy Blood & Tissue Kit (Qiagen) and quality and concentration measured on the NanoDrop OneC spectrophotometer (Thermo Fisher Scientific).

*Reduced Representation Bisulfite Sequencing (RRBS) and identification of differentially methylated regions (DMRs)*

RRBS library was prepared with the Premium RRBS Kit (Diagenode) according to the manufacturer’s instructions with 100 ng of gDNA as starting material. Quality and fragment size were determined with the Bioanalyzer (Agilent). Single read sequencing was performed with a HiSeq2500 machine (51 cycles, Illumina).

The reads were quality- and adapter-trimmed with the Trim Galore! wrapper of cutadapt[11]. The trimmed reads were controlled with FastQC (http://www.bioinformatics.bbsrc.ac.uk/projects/fastqc/). Conversion rates were calculated with custom scripts, counting the amount of G's and C's in non-GC context resulting in values above 99% for all libraries. The reads were mapped to the mm10 version of the mouse genome with BWA[12] and methylCtools[13] after a slightly extended Bis-SNP pipeline[14]. The reads were locally realigned and the quality values were recalibrated before calling the methylation levels. The mm10 SNPs and InDels from dbSNP v138 was used in this process[15]. An initial quality control and exploratory analysis was done with R package RnBeads[16]. Differential loci were detected with MethylKit[17] testing in 500 bp sliding windows with at least 3 CpGs, only including those with a coverage of at least 10x. Differentially methylated regions (DMRs) were defined as ±10% with a *q*-value <0.01.

*Proteomics*

Sample preparation and analysis of mKO mice and WT littermates

Approximately 10 mg of pulverized quadriceps muscle was resuspended in lysis buffer (5% SDS, 10 mM TCEP, 0.1 M TEAB) and lysed by sonication using a PIXUL Multi-Sample Sonicator (Active Motif) with Pulse set to 50, PRF to 1, Process Time to 10 min and Burst Rate to 20 Hz. Lysates were incubated for 10 min at 95°C, alkylated in 20 mM iodoacetamide for 30 min at 25°C and proteins digested using S-Trap™ micro spin columns (Protifi) according to the manufacturer's instructions. Shortly, 12% phosphoric acid was added to each sample (final concentration of phosphoric acid 1.2%) followed by the addition of S-trap buffer (90% methanol, 100 mM TEAB pH 7.1) at a ratio of 6:1. Samples were mixed by vortexing and loaded onto S-trap columns by centrifugation at 4'000 g for 1 min followed by three washes with S-trap buffer. Digestion buffer (50 mM TEAB pH 8.0) containing sequencing-grade modified trypsin (1/25, w/w; Promega) was added to the S-trap column and incubate for 1h at 47°C. Peptides were eluted by the consecutive addition and collection by centrifugation at 4'000 g for 1 min of 40 µl digestion buffer, 40 µl of 0.2% formic acid and finally 35 µl 50% acetonitrile, 0.2% formic acid. Samples were dried under vacuum and stored at -20°C until further use.

Dried peptides were resuspended in 0.1% aqueous formic acid and subjected to LC–MS/MS analysis using a Orbitrap Fusion Lumos Mass Spectrometer fitted with an EASY-nLC 1200 (Thermo Fisher Scientific) and a custom-made column heater set to 60°C. Peptides were resolved using a RP-HPLC column (75 µm × 36 cm) packed in-house with C18 resin (ReproSil-Pur C18–AQ, 1.9 µm resin; Dr. Maisch GmbH) at a flow rate of 0.2 µl/min. The following gradient was used for peptide separation: from 5% B to 12% B over 5 min to 35% B over 65 min to 50% B over 20 min to 95% B over 2 min followed by 18 min at 95% B. Buffer A was 0.1% formic acid in water and buffer B was 80% acetonitrile, 0.1% formic acid in water.

The mass spectrometer was operated in DIA mode. MS1 scans were acquired in the Orbitrap in centroid mode at a resolution of 120'000 FWHM (at 200 m/z), a scan range from 390 to 1'210 m/z, AGC target set to 800 % and a maximum ion injection time of 100 ms. MS2 scans were acquired in the Orbitrap in centroid mode at a resolution of 15'000 FWHM (at 200 m/z), precursor mass range of 400 to 1'200, quadrupole isolation window of 8 m/z with 1 m/z window overlap, a scan range from 145 to 1'450 m/z, AGC target set to Standard and a maximum ion injection time of 22 ms. Peptides were fragmented by HCD (Higher-energy collisional dissociation) with collision energy set to 33% and one microscan was acquired for each spectrum.

The acquired raw-files were searched using the Spectronaut (Biognosys v15.7) directDIA workflow against a murine database (consisting of 17'093 protein sequences downloaded from Uniprot on 20220222) and 392 commonly observed contaminants. Default factory settings were used with minor modifications: In the Pulsar Search Result Filter tab, fragment ion m/z range was set to 300 to 1'800 and the relative intensity minimum was set to 5. Quantitative data was exported from Spectronaut and analyzed using the SafeQuant R package v.2.3.2. https://github.com/eahrne/SafeQuant/)[18]. This analysis included data imputation using the knn algorithm, summation of peak areas per protein and LC-MS/MS run, followed by calculation of protein abundance ratios and testing for differential abundance using empirical Bayes moderated t-statistics (as implemented in the R/Bioconductor limma package). To meet additional assumptions (normality and homoscedasticity) underlying the use of linear regression models and t-tests, MS-intensity signals were transformed from the linear to the log-scale.

Sample preparation and analysis of mTG mice and WT littermates

Quadriceps muscle was pulverized and 10 mg lysed in 8 M Urea, 0.1 M ammonium bicarbonate, phosphatase inhibitors (Sigma) by sonication (Bioruptor, 10 cycles, 30 s on/off, Diagenode) and proteins were digested as described previously[18]. Shortly, proteins were reduced with 5 mM TCEP for 60 min at 37°C and alkylated with 10 mM chloroacetamide for 30 min at 37°C. After diluting samples with 100 mM ammonium bicarbonate buffer to a final urea concentration of 1.6 M, proteins were digested by incubation with sequencing-grade modified trypsin (1/50, w/w; Promega) for 12 h at 37°C. After acidification using 5% TFA, peptides were desalted using C18 reverse-phase spin columns (Macrospin, Harvard Apparatus) according to the manufacturer's instructions, dried under vacuum and stored at -20°C until further use.

Dried peptides were resuspended in 0.1% aqueous formic acid and subjected to LC–MS/MS analysis using a Orbitrap Fusion Lumos Mass Spectrometer fitted with an EASY-nLC 1200 (Thermo Fisher Scientific) and a custom-made column heater set to 60°C. Peptides were resolved using a RP-HPLC column (75 μm × 36 cm) packed in-house with C18 resin (ReproSil-Pur C18–AQ, 1.9 μm resin; Dr. Maisch GmbH) at a flow rate of 0.2 μl/min. The following gradient was used for peptide separation: from 5% B to 12% B over 5 min to 35% B over 65 min to 50% B over 20 min to 95% B over 2 min followed by 18 min at 95% B. Buffer A was 0.1% formic acid in water and buffer B was 80% acetonitrile, 0.1% formic acid in water.

The mass spectrometer was operated in DDA mode with a cycle time of 3 s between master scans. Each master scan was acquired in the Orbitrap at a resolution of 240'000 FWHM (at 200 m/z) and a scan range from 375 to 1'600 m/z followed by MS2 scans of the most intense precursors in the linear ion trap at "Rapid" scan rate with isolation width of the quadrupole set to 1.4 m/z. Maximum ion injection time was set to 50 ms (MS1) and 35 ms (MS2) with an AGC target set to 1e6 and 1e4, respectively. Only peptides with charge state 2 – 5 were included in the analysis. Monoisotopic precursor selection (MIPS) was set to Peptide, and the Intensity Threshold was set to 5e3. Peptides were fragmented by HCD (Higher-energy collisional dissociation) with collision energy set to 35%, and one microscan was acquired for each spectrum. The dynamic exclusion duration was set to 30 s.

The acquired raw-files were imported into the Progenesis QI software (v2.0, Nonlinear Dynamics Limited), which was used to extract peptide precursor ion intensities across all samples applying the default parameters. The generated mgf-file was searched using MASCOT against a murine database (consisting of 17'013 protein sequences downloaded from Uniprot on 20190307) and 392 commonly observed contaminants using the following search criteria: full tryptic specificity was required (cleavage after lysine or arginine residues, unless followed by proline); 3 missed cleavages were allowed; carbamidomethylation (C) was set as fixed modification; oxidation (M) and acetyl (Protein N-term) were applied as variable modifications; mass tolerance of 10 ppm (precursor) and 0.6 Da (fragments). The database search results were filtered to set the false discovery rate (FDR) to 1% on the peptide and protein level, respectively. Quantitative analysis results from label-free quantification were processed using the SafeQuant R package v.2.3.2. (https://github.com/eahrne/SafeQuant/) [18] to obtain peptide relative abundances. This analysis included global data normalization by equalizing the total peak/reporter areas across all LC-MS runs, data imputation using the knn algorithm, summation of peak areas per protein and LC-MS/MS run, followed by calculation of peptide abundance ratios and testing for differential abundance using empirical Bayes moderated t-statistics (as implemented in the R/Bioconductor limma package). To meet additional assumptions (normality and homoscedasticity) underlying the use of linear regression models and t-tests, MS-intensity signals were transformed from the linear to the log-scale.

For all proteomics analyses, only proteins with more than 1 peptide were considered. In addition, a Log2FC cutoff of ±0.2 (proteins with a Log2FC >0.199 or <-0.199) was used and a *p*-value <0.05 was considered statistically significant.

*Single cell and single nucleus RNAseq data analysis*

To create the single-transcriptomic reference dataset including both mononucleated cells and myonuclei, we integrated published single-cell data (scRNAseq) from mononucleated muscle cells[19] and single-nucleus data (snRNAseq) from myonuclei[20]. In detail, we subsetted the provided scRNAseq data from Yang et al. (2022) for all samples from skeletal muscle and reanalyzed it via R/Seurat4.0, including NormalizeData(), FindVariableFeatures() (with top 3000 variable genes), ScaleData() (regressing out mitochondrial genes), and principle component analysis[19]. Then, we integrated the samples using Harmony (github.com/immunogenomics/harmony), applied clustering via FindNeighbors(), FindClusters(), and RunUMAP(). Finally, we annotated clusters based on published marker gene expression and removed cells from high-fat diet-fed mice. To complement the scRNAseq data from mononucleated muscle cells with missing myonuclei data, we used published snRNAseq from tibialis anterior muscles[20]. In short, we applied quality measures as described in the original publication and clustered nuclei with the common Seurat4.0 pipeline. After cluster annotation, we subsetted the dataset for myonuclei only. Subsequent integration of scRNAseq and snRNAseq data was performed by merging all datasets and recalculating the normalization, variable features, scaling, and principle components. We corrected for batch effects and integrated the individual samples via Harmony and clustered as described above. To visualize the expression of a given set of genes, we used the Clustered_DotPlot() function of the "scCustomize" package (samuel-marsh.github.io/scCustomize/) with the minimum color threshold set to zero.

*Statistical analysis*

The statistical analysis of the RNAseq, RRBS and proteomics analysis were done as described in the respective sections. All other statistical analyses were performed in GraphPad Prism 9 using two-tailed Student's t-test. Values are expressed as means ± standard errors of the means (SEM) and generally, a *p*-value <0.05 was considered statistically significant. As an exception, FDR <0.05 and *q*-value <0.01 were considered statistically significant for RNAseq and RRBS analysis, respectively.

### INVENTORY OF SUPPLEMENTAL DATASETS

Table S1. Significantly changed proteins (cutoff: peptide >1; *p*-value <0.05; Log2FC +/-0.2) in quadriceps muscle of sedentary and trained WT and mKO animals as well as sedentary mTG mice. Related to Figures 1, 6, S1, S7.

Table S2: Functional annotation clusters of GO biological processes and REACTOME pathways with an enrichment score >2 using DAVID of significantly changed proteins of trained WT mice as well as sedentary mKO and mTG animals. Related to Figures 1, 6, S1, S7.

Table S3. Functional annotation clusters of GO biological processes and REACTOME pathways with an enrichment score >2 using DAVID of differentially expressed genes in WT mice in response to an acute bout of exercise or after training. Related to Figures 1, 2, S2.

Table S4. All predicted motifs with significantly changed activities using ISMARA of WT and mKO animals in response to an acute bout of exercise or after training. Related to Figures 1, 3, 5, S3, S5, S6.

Table S5. Functional annotation clusters of GO biological processes and REACTOME pathways with an enrichment score >2 using DAVID of proteins that are transcriptionally regulated in either in trained WT muscle of after an acute exercise bout in WT animals. Related to Figure 3.

Table S6. Functional annotation clusters of GO biological processes and REACTOME pathways with an enrichment score >2 using DAVID of differentially expressed genes with differentially methylated regions of sedentary and trained WT and mKO mice as well as sedentary mTG animals. Related to Figures 3, 6, S4, S7.

Table S7. Functional annotation clusters of GO biological processes and REACTOME pathways with an enrichment score >2 using DAVID of differentially expressed genes in mKO mice in response to an acute bout of exercise or after training as well as those that are exclusively regulated in WT muscle. Related to Figures 4, 5, S5, S6.
